# Interacting brains coming in sync through their minds: An inter-brain neurofeedback study

**DOI:** 10.1101/2020.12.16.423070

**Authors:** Viktor Müller, Dionysios Perdikis, Melinda A. Mende, Ulman Lindenberger

**Affiliations:** Center for Lifespan Psychology, Max Planck Institute for Human Development, Berlin, Germany; Brain Simulation Section, Department of Neurology, Charité-Universitätsmedizin Berlin, Germany; Division of Cognitive Sciences, Department of Psychology, University of Potsdam, Potsdam, Germany; Max Planck UCL Centre for Computational Psychiatry and Ageing Research, London, England, and Berlin, Germany

**Keywords:** interpersonal neurofeedback, social interaction, inter-brain synchrony, power spectral density, subjective feeling and appraisal

## Abstract

We live in a social world where we interact with each other. Neurofeedback (NFB) is an indispensable element for such interaction. Single-person NFB studies using electroencephalography (EEG) or other neuroimaging recordings were extensively reported. However, hyperscanning studies using inter-brain synchrony (IBS) as an NFB feature are completely unknown. In this study, we present two different experimental designs where IBS was fed visually back either as two balls approaching each other (so-called *“ball”* design) or as two pendula reflecting oscillatory activity of two participants (so-called *“pendulum”* design). The NFB was provided at two EEG frequencies (2.5 and 5 Hz) and manipulated by enhanced (fake condition) and inverse feedback. We showed that the participants were able to increase IBS by using NFB, especially when it was fed back at the theta frequency. Apart from the intra- and inter-brain coupling, other oscillatory activities (e.g., power spectral density, peak amplitude and peak frequency) changed during the task compared to rest. Moreover, all the measures showed specific correlations to the subjective post-survey item scores, reflecting subjective feeling and appraisal. We conclude that hyperscanning with IBS as a feedback feature seems to be an important tool to examine neural mechanisms of social interaction and collective behavior.

## Introduction

Social interactions are an existential part of human life. Coordinated behavior between two social agents is thought to reflect basic needs and dispositions (Baltes and Staudinger, 1996; Gallese, 2003). Recently, it has been associated with, and linked to, oscillatory couplings between brains (Dumas et al., 2010; Lindenberger et al., 2009). Therefore, investigating both general brain activity within brains and inter-brain coupling has become a topic of research in hyperscanning studies, where two or more participants are recorded simultaneously (see Balconi and Vanutelli, 2017; Czeszumski et al., 2020 for recent reviews). Neurophysiological evidence has shown that brain activity synchronizes in coordinated actions (De Vico Fallani et al., 2010; Dumas et al., 2010; Lindenberger et al., 2009; Müller et al., 2018b, 2013) and bonding behavior(Goldstein et al., 2018; Müller and Lindenberger, 2014). It has also been suggested that the inter-brain coupling in coordinated social interaction is not only caused by similarities in input information and produced output, but reflects, at least in part, neural processes oriented on a temporal adjustment of brain functions and network dynamics (Müller et al., 2018b, 2013; Müller and Lindenberger, 2019; Sänger et al., 2012). However, the neural mechanisms that implement interpersonally coordinated behavior and support social interaction remain elusive and far from understanding (Frith and Frith, 2007; Hari and Kujala, 2009). The present article aims at overcoming limitations and applying the neurofeedback (NFB) methodology in a hyperscanning experiment to test whether the participants are able to synchronize their brains by means of the feedback of neural activity across two brains.

It is well known that humans can train to control their brain activity patterns via NFB (Elmer and Jäncke, 2014; Emmert et al., 2016; Imperatori et al., 2017). NFB may potentially improve the individual’s ability to control brain activity through learned self-regulation when visual or auditory stimuli reflecting electro-cortical activity and excitability are presented or fed back (Birbaumer and Cohen, 2007; Lecomte and Juhel, 2011; Lubar et al., 1995; Rockstroh et al., 1993). Selection of the optimal NFB features is important for success, e.g., feedback type, sensor locations, frequency band, training duration, etc. For example, alpha and theta enhancement training opened new avenues for emotional learning and psychological growth (Chapin and Russell-Chapin, 2014; Demos, 2005) as well as for enhancing mentalization (Imperatori et al., 2017). Even cats can be trained to increase their sensorimotor rhythm (SMR, 12–15 Hz) and via this prevent triggered seizures (Sterman and Friar, 1972). Typically, training lasts over several days or weeks, but also short-term effects are reported. For example, single-channel alpha enhancement with theta crossover can be achieved in less than 30-minute sessions (Demos, 2005). Recently, it has been shown that not only spectral power in different frequency bands, but also coherence and phase synchronization measures between different electrodes, can be used as NFB features (Brunner et al., 2006; Gonuguntla et al., 2013; Gysels and Celka, 2004; Sacchet et al., 2012; Wei et al., 2007). In coherence training, participants are reinforced when correlation or coupling between signals at different brain sites is altered in a desired way. Regarding electrode pairs selection for NFB, two different approaches are used: (i) the random search among all possible electrode pairs and (ii) the selection of appropriate electrode pairs based on physiological prior knowledge (Brunner et al., 2006). Feature selection in an NFB setting can also be achieved using different feature classifiers (Brunner et al., 2006; Eva and Lazar, 2015; Gysels and Celka, 2004; Lotte and Congedo, 2007). A great variety of features has been used for classification, such as amplitude, power or power spectral density (PSD) values, autoregressive (AR) parameters, common special pattern (CSP), and time-frequency features including phase synchronization measures (Brunner et al., 2006; Eva and Lazar, 2015; Gysels and Celka, 2004; Lotte and Congedo, 2007; Wei et al., 2007).

Meanwhile, Brain-Computer-Interfaces (BCI) and NFB technology is not only used for supporting patients (Birbaumer et al., 2000, 1999; Birbaumer and Cohen, 2007; Kotchoubey et al., 1999, 1996; Ramot et al., 2017; Rockstroh et al., 1993; Strehl et al., 2005), but also to improve attention (Jeunet et al., 2020; Linden et al., 1996; Lubar, 1991; Lubar et al., 1995; Lubar and Lubar, 1984; Wang et al., 2011), cognitive performance (Beatty et al., 1974; Rasey et al., 1996; Vernon et al., 2003), and as a means of control in robotics and gaming (Belluomo et al., 2011; Wang et al., 2011). Egner and Gruzelier (2003) also reported an enhancement of music performance under stressful conditions in conservatoire students through NFB training to raise the theta-alpha (5–11 Hz) amplitude. There is also a study on collective NFB in an immersive art environment called “My Virtual Dream” (Kovacevic et al., 2015). The authors reported data from 523 participants collected in a single night, whereby 20 participants at a time experienced a two-part interaction in the dome with an immersive audiovisual environment in front of 50 spectators. They defined two NFB performance measures based on the ability of the participants to maintain the desired state: *relaxation maintenance* obtained by changes in alpha spectral power (alpha performance) and *concentration maintenance* obtained by changes in beta spectral power (beta performance). It has been shown that relaxation conditions showed a gradual decrease of spectral power in mid-range (8–20 Hz) and high frequencies (35–45 Hz), whereas concentration training revealed a gradual increase of power in the beta range and a decrease in low frequencies (<3Hz). The authors indicated that participants were able to learn to modulate their relative spectral power by NFB within only 60 s and 80 s training periods (for relaxation and concentration maintenance, respectively). Whether, and if so, how NFB can influence inter-brain synchronization (IBS) or whether IBS can be used as an NFB feature in a hyper-brain BCI setting is completely unknown. Duan et al. (2013) reported about a cross-brain NFB experiment with two participants, which regulated their near-infrared spectroscopy (NIRS) activity using kinesthetic motor imagery. The difference between the amplitudes of their brain activities at each time point was used as an NFB feature. None of the studies used IBS as an NFB feature to regulate the interaction of the minds. In summary, positive results regarding the intentionally induced or enhanced IBS by means of NFB will provide evidence that IBS is more than just an epiphenomenon of the sensorimotor response and that socially adapted behavior can be altered when a delicate oscillatory balance between the agents is achieved or changed (learned).

The present study was designed to fill the knowledge gap and find out whether the participants can synchronize their brains using NFB. For these purposes, we developed two different NFB experimental designs that reflect inter-brain synchronous electroencephalographic (EEG) activity and feed it visually back: (1) the so-called *“ball”* design reflecting the common inter-brain state of two participants, where the in-phase synchrony between two participants’ fronto-central electrode sites is fed back as two balls approaching each other and (2) the so-called *“pendulum”* design, where the oscillatory activity of two participants (also measured fronto-centrally) is fed back in the form of two pendulums. Thus, in the former case, the participants were able to only control the common inter-brain synchronization state and, in the latter, they were able to control each own oscillatory brain state (pendulum) and adjust both of them to each other.

## Methods

### Participants

All participants had normal or corrected-to-normal vision and did not suffer from any psychological, developmental or neurological disorder. The gender of the participants was balanced so that an equal amount of female-female, female-male, and male-male pairs was tested as it has been found that differences in gender composition of the dyads can influence patterns of inter-brain connectivity (e.g., Baker et al., 2016). Initially, 27 dyads (9 female-female, 9 female-male, 9 male-male) participated in the current study. Data of two of the dyads (1 female-male, 1 male-male) were excluded from the final analysis due to technical issues. Thus, the sample that was included into the final analysis consisted of 25 dyads (9 female-female, 8 female-male, 8 male-male; mean age 26.8 years, SD = 3.1). None of the dyads knew each other before the testing. All participants provided written informed consent prior to the study. The study was approved by the Ethics Committee of Max-Planck Society and was conducted according to the ethical standards which were stated in the Declaration of Helsinki in 1964.

### Psychological Assessment

Immediately before testing, participants were asked to fill in a consumption questionnaire about factors influencing the EEG experiment, for instance consumption of caffeine and alcohol. In addition, educational information was assessed for each participant. After testing, participants filled in two questionnaires while still sitting in the EEG cabin. These were a post-questionnaire about the experiment and a likeability questionnaire (Reysen, 2005). The likeability questionnaire consisted 11 equally pooled items that were transformed to a single likeability scale through averaging across the items. Together with the post-questionnaire, there were overall 16 items reflecting subjective filling, test partner’s likability, and valued capability to influence the task using different strategies, such as concentration, relaxation, thoughts, and mental arithmetic (see Supplementary Table 1 for details). The scores of these items were used for correlation analyses, to provide further information about synchronized states.

### Procedure

During the testing sessions, the dyads were asked to sit back-to-back in an EEG cabin, which was electromagnetically and acoustically shielded. This setup was chosen to prevent participants from communicating with each other or coordinating their movements. Rather, they sat in the EEG cabin and interacted with each other only by knowing that they were performing the NFB task together.

The experiment consisted of 26 sessions. The first session was a pre-rest condition to record a baseline of the brain oscillations of each participant. In this session, participants were asked to sit calmly and relaxed in front of the computer screen without moving their bodies, specifically their limbs, tongue, chin, gums, eyes, and head. The rest condition lasted in total 4 mins. Participants sat 2 mins with opened eyes and another 2 mins with eyes closed. After the relaxation period, the participants performed different NFB tasks: delta and theta *ball* tasks as well as delta and theta *pendulum* tasks. Each task lasted 210 s. The last session was another resting state session and consisted of the same procedure as the resting session at the beginning of the experiment. The participants were asked not to talk to each other throughout the experiment.

### Dyadic Neurofeedback Paradigms

During the NFB tasks, participants were asked to look at the computer screen with an NFB visualization, which consisted of either two balls with different colors (red and blue, respectively) or two pendulums also with the two different colors (red and blue, respectively). They received the instruction to look at the presented feedback and to try to adapt or adjust their brain waves to each other using the NFB. In the *ball* condition, the balls moved toward or away from each other in the horizontal plane in the middle of the screen. When the inter-brain synchronization measured by *ACI* (Absolute Coupling Index) was high, the balls tended to overlap each other; when *ACI* was low, the balls moved to the outer borders of the screen (see Fig. 1 for details). Thus, the task in this condition was to move the balls as close as possible to each other and to make them visually overlapped. In the *pendulum* condition, each of the two pendulums represented the brain oscillations of each of the participants. They were asked to make them swing in-phase, so that pendulum movements were parallel to each other. Thus, in this task, each participant was able to control their own pendulum to achieve in-phase synchrony. See Figure 1 and Supplementary Movies for details.

**Figure 1.**
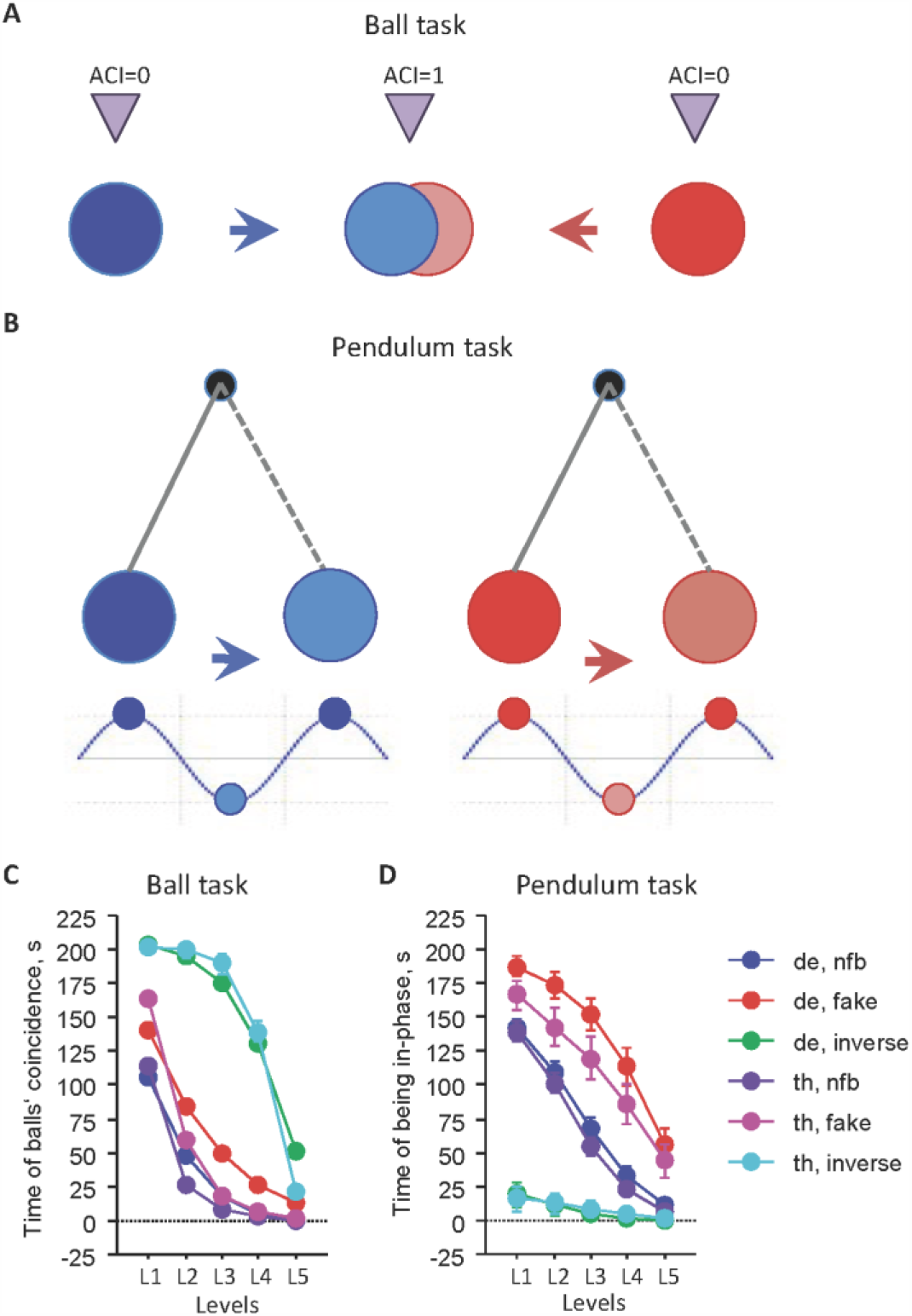
NFB task paradigms and NFB performance. (A) Ball task paradigm. In this task, the balls moved toward or away from each other dependent on the interbrain synchronization measured by the *ACI* at fronto-central electrodes averaged beforehand for each participant’s brain. When the *ACI* is high, the balls tend to overlap each other; when *ACI* is low, the balls move to the outer borders of the screen (B) Pendulum task paradigm. In this task, each of the two pendulums represented the brain oscillations (also at fronto-central sites) of each of the participants. They were asked to make them swing in-phase, so that pendulum movements were parallel to each other. (C) NFB performance in the ball task. The NFB performance in this task was measured as time (in sec) of the balls’ coincidence at five different threshold levels corresponding to synchronization levels measured by *ACI*: L1 (*ACI*>0.5); L2 (*ACI*>0.6); L3 (*ACI*>0.7); L4 (*ACI*>0.8); and L5 (*ACI*>0.9). (D) NFB performance in the pendulum task. The NFB performance in this task was measured as time (in sec) of being in-phase of the two pendula controlled by the two participants. It was also calculated at five different threshold levels dependent on absolute Phase Difference (*aPD*): L1 (*aPD*<0.5π); L2 (*aPD*<0.4π); L3 (*aPD*<0.3π); L4 (*aPD*<0.2π); and L5 (*aPD*<0.1π).

The study contained three different NFB conditions (normal feedback, enhanced feedback, and inverted feedback), which were provided to the dyads in a pseudorandom trial order. In the normal condition, NFB was normally visualized on the screens to give participants an impression of their actual performance. In the trials with enhanced feedback (fake condition), the visualization of NFB was improved to motivate the dyads by giving them the impression that they were performing well in the experiment. In the trials with inverted feedback, the visual feedback improved when the brain oscillations of the dyad were in antiphase or desynchronized (negative learning). These trials were implemented as a control condition to control whether performance was as good as in normal trials. Test subjects were naïve to NFB strategies before the testing. Thus, they were asked to try it using different strategies that came to their minds. Crucially, participants were not triggered by the experimenters toward different strategies before the testing. In addition, participants did not receive any instructions concerning the different experimental conditions (normal feedback, enhanced feedback, inverted feedback) so that they were blind to the testing conditions.

After 210 s, each NFB trial ended indicated by ending the visual feedback on the computer screen. The design of the task was block-wise, meaning that it either started with a block of ball trials or pendulum trials. There was no change of trials inside one block. Within the blocks, there were two sub-blocks based on the calculation of the inter-brain synchronization and corresponding feedback either in the delta (2.5 Hz) or the theta (5.0 Hz) frequency. Within these sub-blocks, there were six trials, always in the same order. The block started with two normal NFB trials, followed by a fake trial with enhanced feedback. The fourth and fifth trials were again normal trials followed by a final inverted trial. This block design led to eight different possible experimental orders which were matched with relation to female-female, female-male, and male-male dyads. In total, each dyad performed 24 trials. Participants took a short break between every five to ten trials so that the duration of the experiment varied between 70 mins and 90 mins.

### EEG Recordings and Offline Analyses

EEG was recorded from the dyads with 64 active Ag/AgCl electrodes per person at a sampling rate of 1000 Hz whereupon the reference electrode was placed at the right mastoid (actiCAP, Brain Products, Munich, Germany). Recorded frequency bands ranged from 0.01 to 250 Hz. The EEG caps were placed on the scalp according to the international 10–10 system. In addition, an electrooculogram in vertical and horizontal dimensions was measured from each participant to control for eye movement. Each participant was recorded with a separate amplifier with separate grounds which were coupled to the same computer. In addition to the EEG measure, heart rate, galvanic skin response, and breathing rate were recorded during the testing. Note that, in this paper, we report only hyperscanning EEG data. For offline analyses, EEG recordings were re-referenced to an average of the left and right mastoid separately for each subject and filtered with a band pass ranging from 0.5 to 100 Hz. The notch filter was set to 50 Hz. Eye-movement correction was accomplished by independent component analysis (Vigário, 1997). For spectral power analyses, the EEG were segmented into 4,096 data-points epochs with 50% overlap. The power spectral density (PSD) was calculated using Welch’s method and Hanning window function in the four frequency ranges: delta (0.5–4 Hz), theta (4–8 Hz), alpha (8–14 Hz), and beta (14–28 Hz). Within these frequency ranges, the peak amplitude and peak frequency were determined. Intra- and inter-brain synchronization were assessed using the *ACI* measure (Müller et al., 2013; Müller and Lindenberger, 2011) for all possible electrode pairs within and between the brains, respectively. The *ACI* reflecting in-phase synchronization was determined for four frequency bins (2.5, 5, 10, and 20 Hz) within the epochs of 5 s without overlap. For each electrode location, *ACI* coupling strengths were calculated as a sum of all coupling pairs from one electrode to all other separately for within- and between-brain coupling. For statistical analyses, all the measures were averaged across the epochs.

### Measure of Neurofeedback Performance

The NFB was calculated online using six fronto-central channels from each participant (F3, Fz, F4, C3, Cz, and C4), which were averaged within the two brains, to create a single feedback time series for each participant (*i*):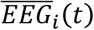. These time series were determined for the time segment of 4 s centered around time *t*. The complex Gabor wavelet time-frequency transform was computed on 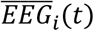, in order to extract the unwrapped instantaneous phase of each participant *φ*_*i*_(*t*), which was used in a different way for each of the two experimental designs (see below). The Gabor wavelet was computed with cycle number *c* = 10 following the equation (Lachaux et al., 1999):

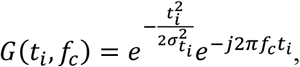

where *t*_*i*_ and *f*_*c*_ denote time and the center frequency, respectively, 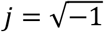, and 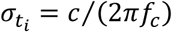, leading to a time wavelet length of 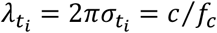 and to a spectral bandwidth of 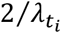 Hz. The *f*_*c*_ was equal to 2.5 Hz for the delta and 5 Hz for the theta band, leading to a time wavelet length of 4 s and 2 s, respectively.

### Ball paradigm

The instantaneous phase difference was computed, wrapped in the [− π, +π] interval:

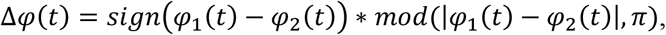

here *sign*(·) is the function that returns the sign of its argument, |·| denotes the absolute value, and *mod*(·)is the modulus function that returns the remainder of the division of its arguments. We used the instantaneous phase difference to calculate the *ACI* (Müller et al., 2013; Müller and Lindenberger, 2011) within the 4 s segment. *ACI* computes the ratio of time points of the phase difference lying within the interval [− *π*/4, + *π*/4] to the total number of points in the whole interval, that is, it evaluates the most-strict sense of in-phase synchronization as a stable and small phase difference or time coincidence of two oscillations.

Then, *ACI*(*t*) was law pass filtered via a linear difference equation, in order to get the final feedback signal:

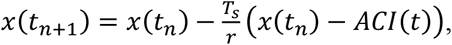

where *t*_*n*_ is the n^th^ time segment, *T*_*s*_ = 17 ms denotes the time resolution of the feedback, and *r* = 15*T*_*s*_ is a time scale factor. The initial condition was set to *x*(1) = *ACI*(1). Thus, the feedback followed *ACI*(*t*) asymptotically and smoothly and ranged in the interval [0, 1].

A quantity computed as *y*(*t*_*n*_) = (1 + *IMF*)*x*(*t*_*n*_), where *IMF* was the “improvement factor”, set to 0.3 for “enhanced” trials and 0.0 for “normal” ones, guided the positions of the two balls, which were drawn at the display of the experimental stimulus computer. Note that *y*(*t*_*n*_) = 0 corresponded to the maximum distance of the screen and *y*(*t*_*n*_) ≥ 1 to fully overlapping balls. For “inverted” trials, this correspondence was the opposite.

### Pendulum Paradigm

Since the delta and theta frequencies that we used to compute the feedback features are too fast for the visualization of the pendulum, we used a “Cross-Frequency Coupling factor” (*CFCf*) to transform the oscillations at these frequencies to the low-frequency oscillations of the pendulum (about 0.25 Hz). The *CFCf* was set to 10 for the delta and 20 for the theta frequency, leading to 1 full cycle of the pendulum feedback for each 10 (20) cycles of the delta (theta) band EEG feedback input, respectively (i.e., the frequency of the pendulum feedback was always fixed to 0.25 Hz for both frequencies). For these purposes, the unwrapped phases of the last time point of the buffer were divided by *CFCf* and then used to set the position of each pendulum. The latter was scaled so that the angular position during a full oscillation cycle of each pendulum varied in the interval [− *π*/6, + *π*/6].

For the “enhanced” condition, a modification to the pendula angular positions was made, such that their angular distance was improved by an *improvement factor* (*IMF* = 30°):

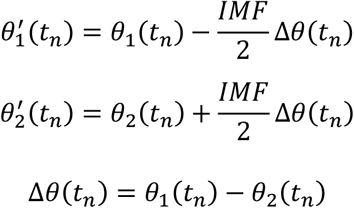

where *θ*_1,2_(*t*_*n*_) and 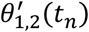 are the angular positions before and after the correction, respectively.

Furthermore, the initial angular positions of the pendulums for “normal trials” differed by an angular factor of 60°, in order to prevent the initial feedback from looking too good, which would be the case if the pendulums started in parallel. Instead for the “enhanced” trials, the initial angular factor was only 30° so that the feedback looked better than in “normal” trials. In “inverted” trials, no initial angular factor was used.

### EEG Real Time Feedback Generation

The feedback EEG signal was sent in packages of *T*_*s*_ = 17 ms, which was the time resolution of the feedback. The packages were gathered into a buffer of 4 s and we further processed such time segments (i.e., of the last 4 s) to generate the feedback signal. The display was refreshed approximately for every new feedback data package, since its refresh rate was 60 Hz, resulting in a refresh time of 17 ms.

The whole experiment was performed in MATLAB 2012b (The MathWorks, Inc., Natick, Massachusetts, United States) code (including the statistics’ toolbox and psychtoolbox), written by the authors. For the real-time connection of the recording computer to the experimental stimulus computer and vice versa, we have used scripts provided by Brain Products GmbH (Gilching, Germany).

## Statistical analysis

For statistical analyses, the EEG spectral measures (PSD, peak amplitude and peak frequency) and *ACI* coupling strengths (sum of all coupling pairs from one electrode to all other) were first determined for the 60 electrode locations and then collapsed into 9 brain sites: FL (frontal left), FZ (mid frontal), FR (frontal right), CL (central left), CZ (mid central), CR (central right), PL (parietal left), PZ (mid parietal), and PR (parietal right). All the measures were analyzed using three-way repeated measures ANOVAs with within-subject factors *Condition* (rest, nfb, fake, and inverse task conditions), *Antero-Posterior* (F=frontal, C=central, and P=parietal), and *Medio-Lateral* (L=left, Z=mid, and R=right). To investigate how stable the peak frequencies were within the pairs, we determined the peak frequency difference changes within the test pairs and subjected them to the ANOVA as before. Greenhouse-Geisser epsilons were used in all ANOVAs for a non-sphericity correction when necessary. The Student-Newman-Keuls (SNK) test was employed for *post-hoc* testing. To relate the EEG indices to behavioral measures, we correlated the former with the 16 items of the post-survey, reflecting subjective filling, test partner’s likability, and valued capability, to influence the task. The Spearman’s rank correlation was used for this purpose.

## Results

### Neurofeedback Performance

To better understand changes in EEG dynamics, we firstly present NFB task performance as this was shown during the experiment on the screen. Note that this performance in fake and inverse task conditions results from the corresponding manipulations as described in Methods. In other words, this performance indicates results of the feedback applied during the experiment. Examples of different task conditions are shown in Supplementary Movies. The task performance was measured as the time of the balls’ coincidence in the ball task and as the time of being in-phase in the pendulum task. The NFB task performance was calculated at five different threshold levels. The threshold levels indicated synchronization levels measured by *ACI* (ball task) or phase difference (pendulum task). In the former case, the threshold levels correspond to synchronization levels of: L1 (*ACI*>0.5); L2 (*ACI*>0.6); L3 (*ACI*>0.7); L4 (*ACI*>0.8); and L5 (*ACI*>0.9). In the latter case, the threshold level was calculated dependent on absolute phase difference (*aPD*): L1 (*aPD*<0.5π); L2 (*aPD*<0.4π); L3 (*aPD*<0.3π); L4 (*aPD*<0.2π); and L5 (*aPD*<0.1π). Results of this analysis are presented in Figures 1C and 1D. It can be seen that performance declined exponentially with the increasing synchronization threshold. In the ball task, the highest performance (time of balls’ coincidence in sec) was in the inverse task condition because the lowest synchronization was transformed in this task condition to its inverse, that is, the balls overlapped for most of the time (high coincidence time). This corresponds then to the negative learning. The next best performance was achieved during the fake condition, because the performance shown during the task (real feedback) was improved by the improvement factor (see Methods for details). The lowest performance was achieved during the normal NFB task condition. In the pendulum task, the performance (time of being in-phase in sec) was the highest in the fake task condition, also due to the improvement of the feedback through the improvement factor, and the lowest in the inverse task condition. The latter probably happened because the movement of the pendulums always began in anti-phase and it was difficult for the participants to get them in sync. Contrary to the ball task with negative learning in the inverse condition, in the pendulum task, we have experience a negative feedback frustrating the participants through an apparent inefficiency to control the pendula or to improve their performance.

### Power Spectral Density (PSD)

Before describing phase synchronization changes under different NFB conditions, we first present PSD changes to show how, overall, the oscillatory brain activity is evolving or changing under these conditions. For these purposes, we not only concentrate on the frequencies of interest (FOI) manipulated during the task (i.e., delta and theta), but also will describe other frequencies (e.g., alpha and beta) to better understand the effects.

Corresponding log-transformed PSD changes were analyzed using three-way repeated measures ANOVAs (*Condition* × *Antero-Posterior* × *Medio-Lateral*). Results of these ANOVAs for the four frequency bands restricted to the factor *Condition* and interactions with this factor are presented in Table 1 and Figure 2. It can be seen that the main effect *Condition* and the interaction *Condition* × *Antero-Posterior* were significant practically in all four frequency bands, with the exception of the *Condition* × *Antero-Posterior* interaction in the delta pendulum task (see Table 1 for details). The SNK post hoc test revealed a significant decrease of PSD in the NFB tasks compared to the rest condition in all four frequency bands (see Fig. 2 for details). This decrease of PSD in task conditions compared with rest takes place to a greater extent at parietal electrode sites. The significant interaction *Condition* × *Medio-Lateral* in the delta and theta ball task indicates a stronger decrease of the theta PSD at the left and right lateral sites compared to the medial sites.

**Table 1:** ANOVA results for the PSD across the four frequency bands in the four NFB tasks

| Design | Ball design |  |  |  |  |  | Pendulum design |  |  |  |  |  |
| --- | --- | --- | --- | --- | --- | --- | --- | --- | --- | --- | --- | --- |
| Task | Delta |  |  | Theta |  |  | Delta |  |  | Theta |  |  |
| Value | F | P | Eta<sup>2</sup> | F | P | Eta<sup>2</sup> | F | P | Eta<sup>2</sup> | F | P | Eta<sup>2</sup> |
| Delta PSD |  |  |  |  |  |  |  |  |  |  |  |  |
| C | 5.99 | 0.004 | 0.11 | 6.45 | 0.004 | 0.12 | 4.35 | 0.018 | 0.082 | 4.18 | 0.019 | 0.079 |
| C × AP | 2.88 | 0.042 | 0.056 | 2.70 | 0.043 | 0.052 | 2.27 | 0.077 | 0.044 | 2.33 | 0.075 | 0.045 |
| C × ML | 1.70 | 0.15 | 0.034 | 1.34 | 0.25 | 0.027 | 0.63 | 0.67 | 0.013 | 0.94 | 0.45 | 0.019 |
| C × AP × ML | 0.69 | 0.69 | 0.014 | 0.76 | 0.61 | 0.015 | 1.42 | 0.20 | 0.028 | 3.09 | 0.003 | 0.059 |
| Theta PSD |  |  |  |  |  |  |  |  |  |  |  |  |
| C | 26.86 | 0.000 | 0.35 | 25.95 | 0.000 | 0.35 | 25.57 | 0.000 | 0.34 | 24.27 | 0.000 | 0.33 |
| C × AP | 6.71 | 0.000 | 0.12 | 7.56 | 0.000 | 0.13 | 3.78 | 0.011 | 0.072 | 5.82 | 0.001 | 0.11 |
| C × ML | 3.22 | 0.015 | 0.062 | 2.45 | 0.047 | 0.048 | 0.92 | 0.46 | 0.018 | 1.40 | 0.24 | 0.028 |
| C × AP × ML | 0.58 | 0.77 | 0.012 | 0.56 | 0.75 | 0.011 | 0.86 | 0.51 | 0.017 | 3.23 | 0.002 | 0.062 |
| Alpha PSD |  |  |  |  |  |  |  |  |  |  |  |  |
| C | 100.66 | 0.000 | 0.67 | 89.54 | 0.000 | 0.65 | 72.82 | 0.000 | 0.60 | 67.22 | 0.000 | 0.58 |
| C × AP | 15.99 | 0.000 | 0.25 | 18.94 | 0.000 | 0.28 | 15.51 | 0.000 | 0.24 | 15.46 | 0.000 | 0.24 |
| C × ML | 1.00 | 0.41 | 0.020 | 0.37 | 0.81 | 0.008 | 1.38 | 0.24 | 0.027 | 0.28 | 0.89 | 0.006 |
| C × AP × ML | 1.25 | 0.28 | 0.025 | 0.56 | 0.75 | 0.011 | 0.50 | 0.77 | 0.010 | 1.72 | 0.11 | 0.034 |
| Beta PSD |  |  |  |  |  |  |  |  |  |  |  |  |
| C | 22.06 | 0.000 | 0.31 | 27.68 | 0.000 | 0.36 | 20.07 | 0.000 | 0.29 | 19.71 | 0.000 | 0.29 |
| C × AP | 8.28 | 0.000 | 0.15 | 8.98 | 0.000 | 0.16 | 10.68 | 0.000 | 0.18 | 8.94 | 0.001 | 0.15 |
| C × ML | 1.07 | 0.38 | 0.021 | 0.34 | 0.87 | 0.007 | 1.62 | 0.16 | 0.032 | 0.52 | 0.76 | 0.010 |
| C × AP × ML | 0.61 | 0.76 | 0.012 | 1.42 | 0.21 | 0.028 | 1.14 | 0.34 | 0.023 | 2.37 | 0.018 | 0.046 |

C=Condition; AP=Antero-Posterior; ML=Medio-Lateral

**Figure 2.**
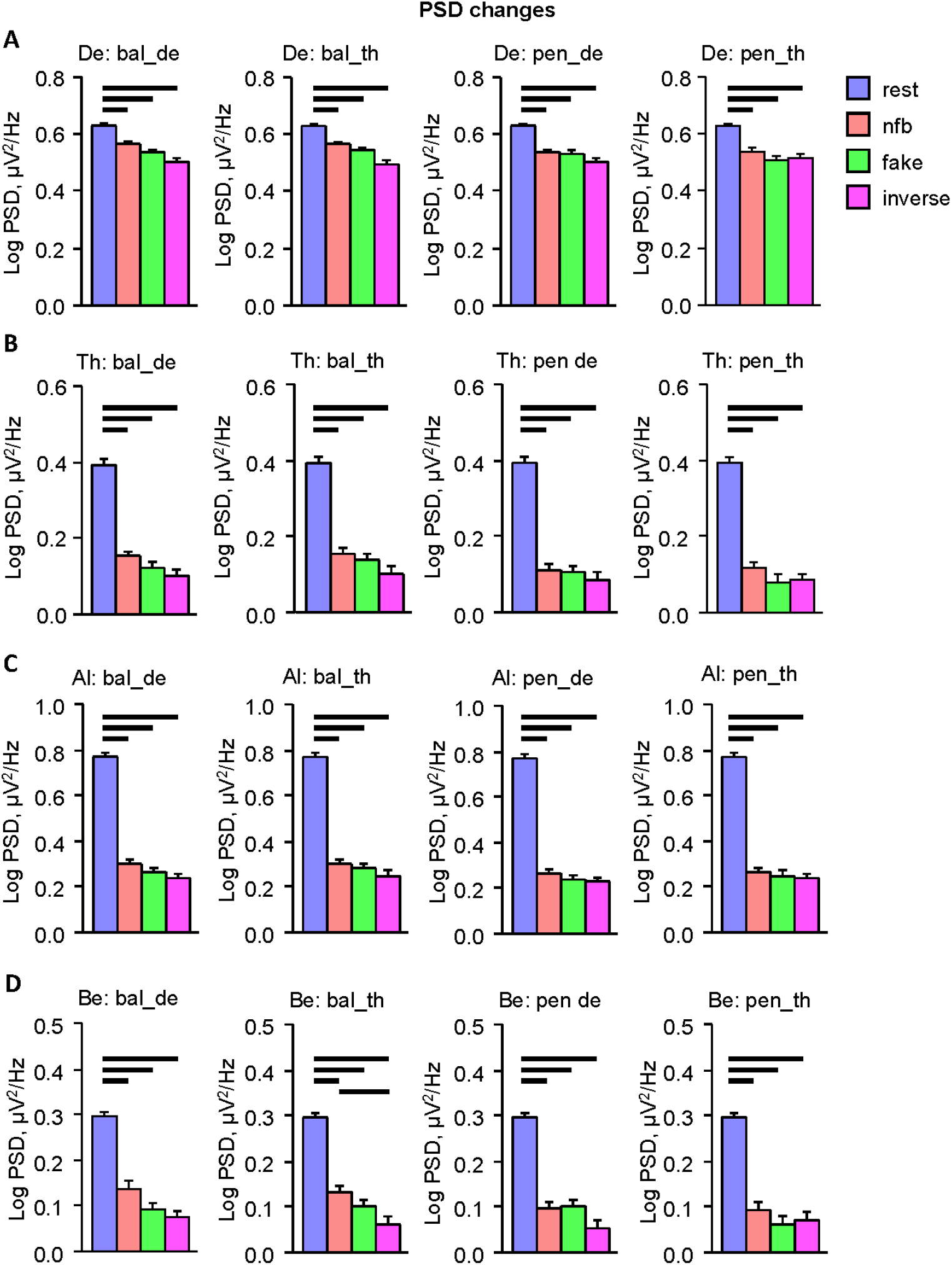
PSD changes across the task conditions in the four frequency bands. (A) Delta PSD across the four NFB tasks and the four task conditions. (B) Theta PSD across the four NFB tasks and the four task conditions. (C) Alpha PSD across the four NFB tasks and the four task conditions. (D) Beta PSD across the four NFB tasks and the four task conditions. Bar diagrams represent PSD means and standard errors across the four task conditions: resting state, normal NFB, fake, and inverse task conditions. NFB tasks: Ball delta task = bal_de; ball theta task = bal_th; pendulum delta task = pen_de; pendulum theta task = pen_th. Horizontal bold lines indicate significant differences as revealed by the SNK post-hoc test (*p* < 0.05).

Further, we correlated PSD during the NFB task condition with 16 post-survey item scores reflecting subjective feeling, test partner’s likability, and valued capability to influence the task (see Supplementary Table S1 for details). The results of these correlation analyses are presented in Figure 3. There were relatively strong positive correlations of PSD in theta, alpha, and beta frequency bands with tiredness at the end of the experiment (item 2) and nervousness (item 3); delta PSD correlated positively with tiredness at the end of the experiment and expectation of an important appointment (item 4). Note that there were no significant correlations with tiredness at the beginning of the experiment (item 1). In all frequency bands, PSD correlated negatively with the sympathy for the test partner (item 8). In addition, there were positive correlations between delta PSD (during the ball task) and valued capability to influence the task through relaxation (item 14) as well as also negative correlations of theta, alpha, and beta PSD with some other scores of the valued capability to influence the task (see Figure 3 for details).

**Figure 3.**
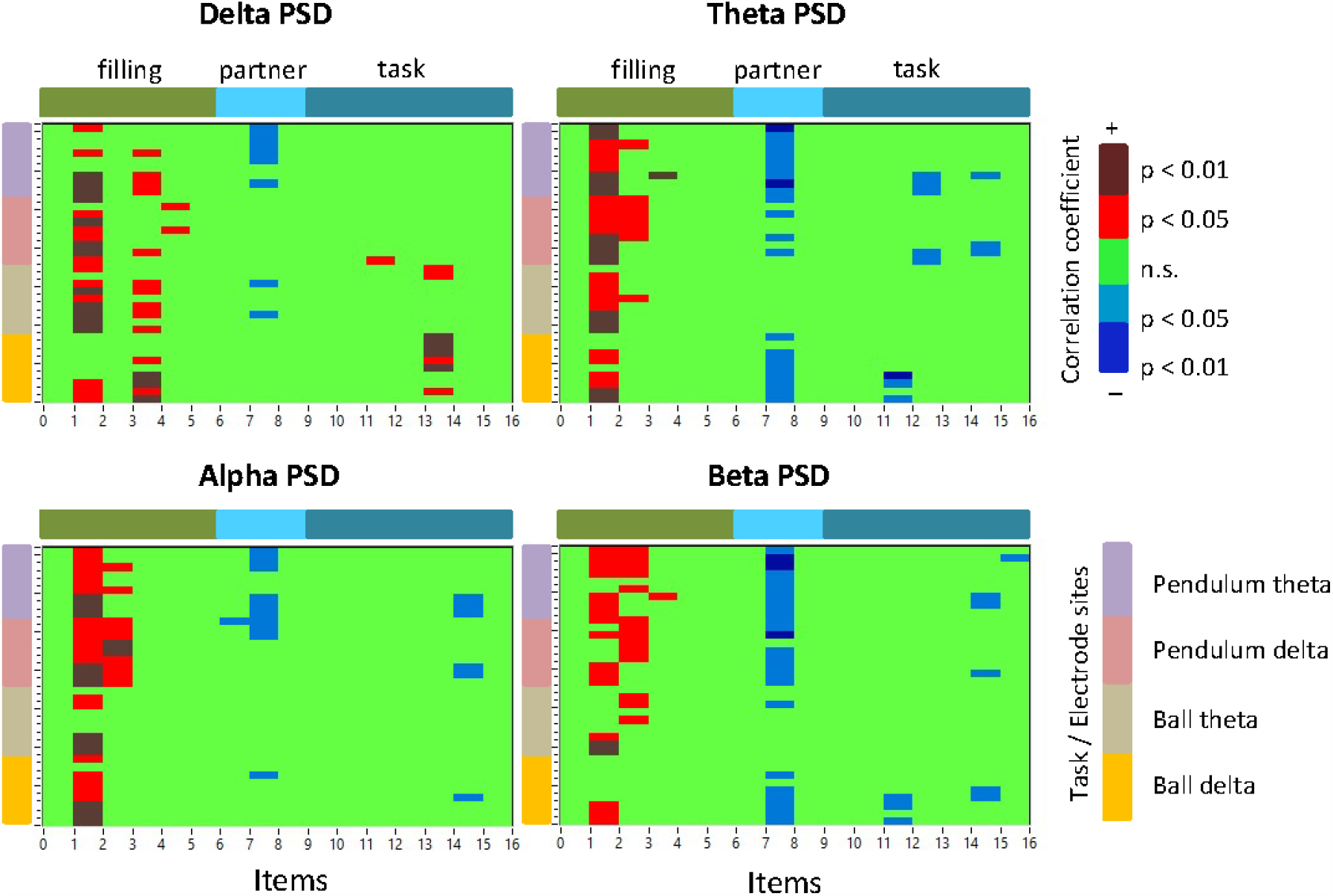
Correlations between the 16 post-survey item scores and PSD in the four frequency bands. The x-axis represents the 16 post-survey item scores, which are divided into three subgroups (filing-, partner-, and task-related) as indicated by the color bars at the top of the diagrams. The y-axis represents the four tasks with nine electrode sites for each (see legend Task/Electrode sites). Electrode sites are ordered as follows: FL (frontal left), FZ (mid frontal), FR (frontal right), CL (central left), CZ (mid central), CR (central right), PL (parietal left), PZ (mid parietal), and PR (parietal right). The diagrams show only significant (negative and positive) correlation coefficients as indicated in the legend.

### Power Spectral Peak Amplitude and Peak Frequency

We further investigated the power spectral peak amplitude and peak frequency during the task and rest conditions. Whereas the peak amplitude changes were relatively similar to the PSD changes (see Supplementary Table S2 and Fig. S1 for details), the peak frequency revealed a significant main effect of *Condition* in the delta frequency band during the pendulum tasks and in the theta frequency band among all NFB tasks. The alpha peak frequency showed a significant interaction *Condition* × *Medio-Lateral* among all tasks and a significant interaction *Condition* × *Antero-Posterior* × *Medio-Lateral* during the ball tasks. The beta peak frequency revealed a significant interaction *Condition* × *Antero-Posterior* among all the NFB tasks and a significant interaction *Condition* × *Medio-Lateral* during the delta ball and delta pendulum tasks as well as a significant interaction *Condition* × *Antero-Posterior* × *Medio-Lateral* during the last task (see Table 2 for details). As shown in Figure 4, the delta (in the pendulum tasks) and the theta peak frequencies decrease during the NFB tasks compared to the rest condition. Whereas the delta peak frequency, distributed around 1 Hz, was far from the delta task frequency (i.e., 2.5 Hz), the theta peak frequency approached from about 6 Hz during the rest condition to the frequency of about 5.5 Hz during the task conditions, which is relatively close to the theta task frequency (i.e., 5 Hz). Interestingly, the alpha peak frequency diverges in the lateral (left and right) sites compared to the medial sites, whereby the alpha peak frequency at the lateral sites increases in task conditions (especially in NFB and fake conditions) compared to the rest and remains more stable at medial sites (see Fig. 4 for details). The beta peak frequency also diverges from the rest condition to the task conditions in antero-posterior sites, whereby the beta peak frequency increases at central, and especially frontal sites, and decreases at the parietal sites (see Fig. 4 for details). In general, we can see the facilitation of high alpha and beta peak frequencies in the task conditions compared to the rest, at least in several brain regions.

**Table 2:**
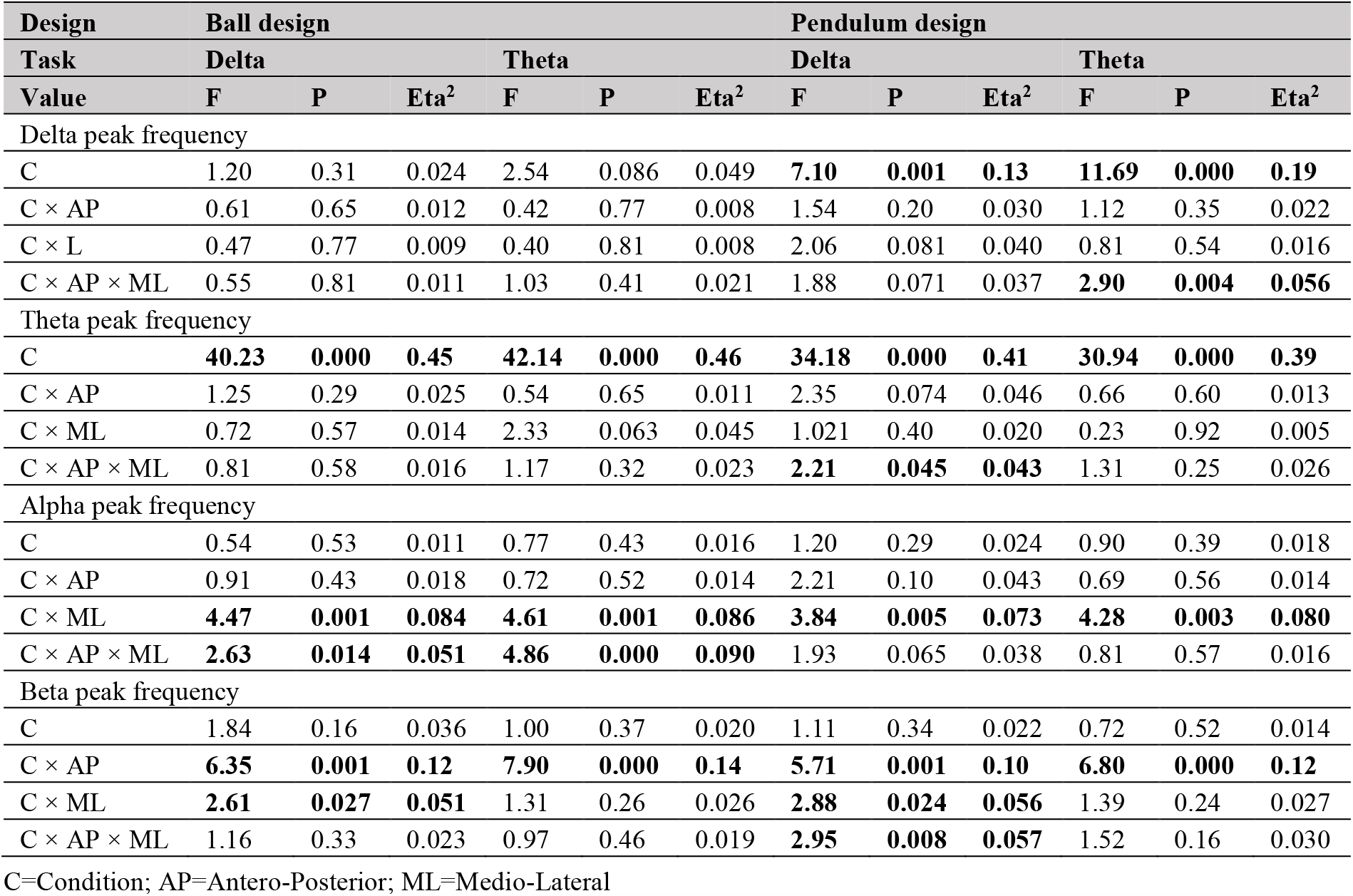
ANOVA results for the peak frequency across the four frequency bands in the four NFB tasks

| Design | Ball design |  |  |  |  |  | Pendulum design |  |  |  |  |  |
| --- | --- | --- | --- | --- | --- | --- | --- | --- | --- | --- | --- | --- |
| Task | Delta |  |  | Theta |  |  | Delta |  |  | Theta |  |  |
| Value | F | P | Eta<sup>2</sup> | F | P | Eta<sup>2</sup> | F | P | Eta<sup>2</sup> | F | P | Eta<sup>2</sup> |
| Delta peak frequency |  |  |  |  |  |  |  |  |  |  |  |  |
| C | 1.20 | 0.31 | 0.024 | 2.54 | 0.086 | 0.049 | **7.10** | **0.001** | **0.13** | **11.69** | **0.000** | **0.19** |
| C × AP | 0.61 | 0.65 | 0.012 | 0.42 | 0.77 | 0.008 | 1.54 | 0.20 | 0.030 | 1.12 | 0.35 | 0.022 |
| C × L | 0.47 | 0.77 | 0.009 | 0.40 | 0.81 | 0.008 | 2.06 | 0.081 | 0.040 | 0.81 | 0.54 | 0.016 |
| C × AP × ML | 0.55 | 0.81 | 0.011 | 1.03 | 0.41 | 0.021 | 1.88 | 0.071 | 0.037 | **2.90** | **0.004** | **0.056** |
| Theta peak frequency |  |  |  |  |  |  |  |  |  |  |  |  |
| C | **40.23** | **0.000** | **0.45** | **42.14** | **0.000** | **0.46** | **34.18** | **0.000** | **0.41** | **30.94** | **0.000** | **0.39** |
| C × AP | 1.25 | 0.29 | 0.025 | 0.54 | 0.65 | 0.011 | 2.35 | 0.074 | 0.046 | 0.66 | 0.60 | 0.013 |
| C × ML | 0.72 | 0.57 | 0.014 | 2.33 | 0.063 | 0.045 | 1.021 | 0.40 | 0.020 | 0.23 | 0.92 | 0.005 |
| C × AP × ML | 0.81 | 0.58 | 0.016 | 1.17 | 0.32 | 0.023 | **2.21** | **0.045** | **0.043** | 1.31 | 0.25 | 0.026 |
| Alpha peak frequency |  |  |  |  |  |  |  |  |  |  |  |  |
| C | 0.54 | 0.53 | 0.011 | 0.77 | 0.43 | 0.016 | 1.20 | 0.29 | 0.024 | 0.90 | 0.39 | 0.018 |
| C × AP | 0.91 | 0.43 | 0.018 | 0.72 | 0.52 | 0.014 | 2.21 | 0.10 | 0.043 | 0.69 | 0.56 | 0.014 |
| C × ML | **4.47** | **0.001** | **0.084** | **4.61** | **0.001** | **0.086** | **3.84** | **0.005** | **0.073** | **4.28** | **0.003** | **0.080** |
| C × AP × ML | **2.63** | **0.014** | **0.051** | **4.86** | **0.000** | **0.090** | 1.93 | 0.065 | 0.038 | 0.81 | 0.57 | 0.016 |
| Beta peak frequency |  |  |  |  |  |  |  |  |  |  |  |  |
| C | 1.84 | 0.16 | 0.036 | 1.00 | 0.37 | 0.020 | 1.11 | 0.34 | 0.022 | 0.72 | 0.52 | 0.014 |
| C × AP | **6.35** | **0.001** | **0.12** | **7.90** | **0.000** | **0.14** | **5.71** | **0.001** | **0.10** | **6.80** | **0.000** | **0.12** |
| C × ML | **2.61** | **0.027** | **0.051** | 1.31 | 0.26 | 0.026 | **2.88** | **0.024** | **0.056** | 1.39 | 0.24 | 0.027 |
| C × AP × ML | 1.16 | 0.33 | 0.023 | 0.97 | 0.46 | 0.019 | **2.95** | **0.008** | **0.057** | 1.52 | 0.16 | 0.030 |

C=Condition; AP=Antero-Posterior; ML=Medio-Lateral

**Figure 4.**
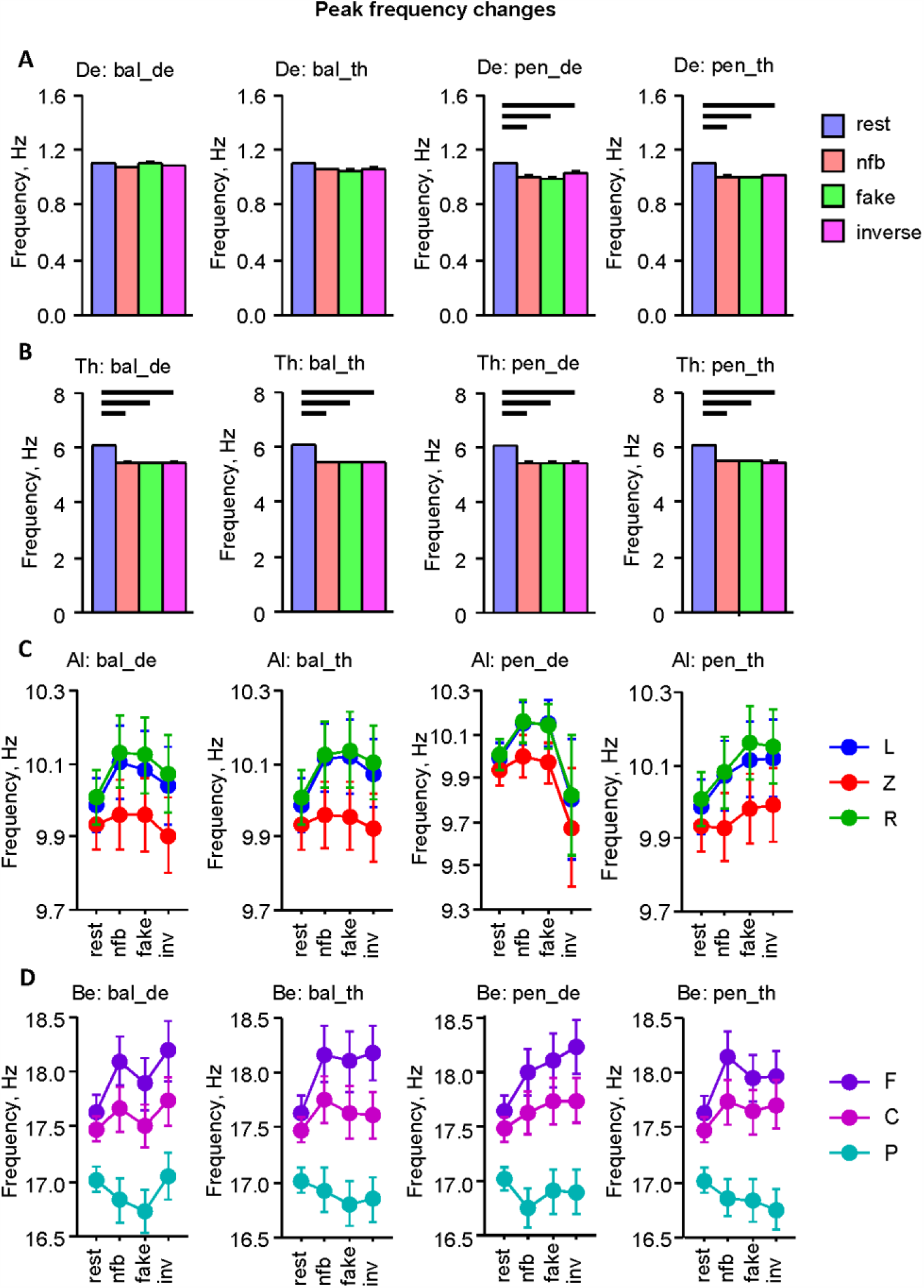
Peak frequency changes across the task conditions in the four frequency bands. (A) Delta peak frequency across the four NFB tasks and the four task conditions. (B) Theta peak frequency across the four NFB tasks and the four task conditions. (C) Alpha peak frequency across the four NFB tasks and the four task conditions at the mid-lateral sites. (D) Beta PSD across the four NFB tasks and the four task conditions at the fronto-parietal sites. Bar diagrams represent PSD means and standard errors across the four task conditions: resting state, normal NFB, fake, and inverse task conditions. Line diagrams represent the *Condition* by *Medio-Lateral* or *Condition* by *Antero-Posterior* interactions. NFB tasks: Ball delta task = bal_de; ball theta task = bal_th; pendulum delta task = pen_de; pendulum theta task = pen_th. Medio-lateral sites: L = left; Z = medio; R = right. Antero-posterior sites: F = frontal; C = central; P = parietal. Horizontal bold lines indicate significant differences as revealed by the SNK post-hoc test (*p* < 0.05).

Most interestingly, we analyzed the peak frequency differences within the pairs. The results of these analyses are presented in Table 3 and Figure 5. It can be seen that the task frequencies, i.e., delta and theta peak frequencies, did not show any significant changes across conditions, whereas alpha and beta peak frequencies revealed a significant main effect of the factor *Condition* in all four tasks, with the exception of non-significant differences in the alpha peak frequency during the delta ball task. As shown in Figure 5, both the alpha and the beta peak frequency differences within the participant pairs increase in the task conditions compared to rest. No significant differences were found between the NFB task conditions themselves.

**Table 3:** ANOVA results for the peak frequency differences within the test pairs across the four frequency bands in the four NFB tasks

| Design | Ball design |  |  |  |  |  | Pendulum design |  |  |  |  |  |
| --- | --- | --- | --- | --- | --- | --- | --- | --- | --- | --- | --- | --- |
| Task | Delta |  |  | Theta |  |  | Delta |  |  | Theta |  |  |
| Value | F | P | Eta<sup>2</sup> | F | P | Eta<sup>2</sup> | F | P | Eta<sup>2</sup> | F | P | Eta<sup>2</sup> |
| Delta peak frequency difference |  |  |  |  |  |  |  |  |  |  |  |  |
| C | 1.71 | 0.17 | 0.067 | 0.72 | 0.54 | 0.029 | 1.14 | 0.33 | 0.045 | 2.16 | 0.12 | 0.083 |
| C × AP | 0.40 | 0.82 | 0.016 | 0.78 | 0.56 | 0.031 | 0.20 | 0.95 | 0.008 | 0.85 | 0.49 | 0.034 |
| C × L | 1.21 | 0.31 | 0.048 | 2.30 | 0.061 | 0.087 | 1.68 | 0.16 | 0.065 | 2.08 | 0.080 | 0.080 |
| C × AP × ML | 1.12 | 0.35 | 0.045 | 0.97 | 0.46 | 0.039 | 0.59 | 0.75 | 0.024 | 1.18 | 0.32 | 0.047 |
| Theta peak frequency difference |  |  |  |  |  |  |  |  |  |  |  |  |
| C | 0.72 | 0.54 | 0.029 | 0.20 | 0.73 | 0.008 | 0.19 | 0.75 | 0.008 | 0.25 | 0.86 | 0.010 |
| C × AP | 0.42 | 0.78 | 0.017 | 0.72 | 0.57 | 0.029 | 0.66 | 0.64 | 0.027 | 1.73 | 0.15 | 0.067 |
| C × ML | 0.25 | 0.88 | 0.010 | 0.49 | 0.77 | 0.020 | 0.98 | 0.42 | 0.039 | 0.85 | 0.49 | 0.034 |
| C × AP × ML | 0.72 | 0.64 | 0.029 | 0.35 | 0.93 | 0.014 | 1.72 | 0.11 | 0.067 | 1.44 | 0.20 | 0.057 |
| Alpha peak frequency difference |  |  |  |  |  |  |  |  |  |  |  |  |
| C | 1.88 | 0.14 | 0.073 | **4.04** | **0.030** | **0.14** | **3.13** | **0.041** | **0.12** | **5.30** | **0.006** | **0.18** |
| C × AP | 0.41 | 0.78 | 0.017 | 0.37 | 0.82 | 0.015 | 1.47 | 0.22 | 0.058 | 0.13 | 0.97 | 0.005 |
| C × ML | 0.60 | 0.65 | 0.024 | 1.56 | 0.19 | 0.061 | 2.34 | 0.056 | 0.089 | 0.27 | 0.91 | 0.011 |
| C × AP × ML | 0.62 | 0.72 | 0.025 | 0.47 | 0.85 | 0.019 | 1.35 | 0.23 | 0.053 | 0.67 | 0.67 | 0.027 |
| Beta peak frequency difference |  |  |  |  |  |  |  |  |  |  |  |  |
| C | **4.91** | **0.008** | **0.17** | **8.47** | **0.001** | **0.26** | **6.97** | **0.001** | **0.23** | **7.26** | **0.001** | **0.23** |
| C × AP | 0.46 | 0.77 | 0.019 | 1.22 | 0.31 | 0.048 | 0.73 | 0.58 | 0.030 | 0.52 | 0.72 | 0.021 |
| C × ML | 1.59 | 0.17 | 0.062 | 1.88 | 0.12 | 0.073 | **3.61** | **0.008** | **0.13** | 2.30 | 0.058 | 0.088 |
| C × AP × ML | 1.13 | 0.35 | 0.045 | 0.73 | 0.63 | 0.030 | 0.49 | 0.80 | 0.020 | 0.73 | 0.63 | 0.030 |

C=Condition; AP=Antero-Posterior; ML=Medio-Lateral

**Figure 5.**
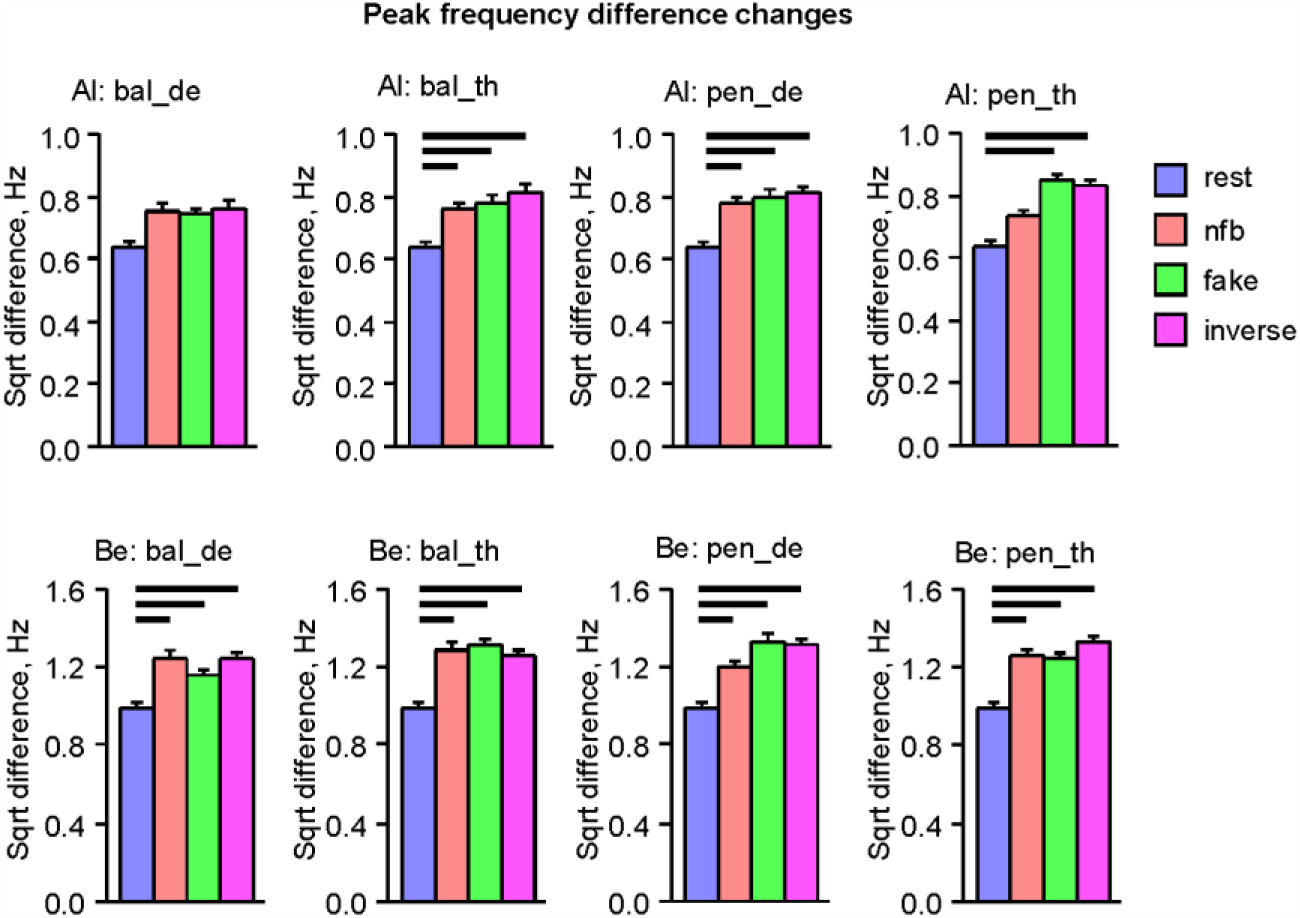
Peak frequency difference changes within the test pairs across the task conditions in the alpha and beta frequency bands. (A) Alpha peak frequency differences within the test pairs across the four NFB tasks and the four task conditions. (B) Beta peak frequency differences within the test pairs across the four NFB tasks and the four task conditions. NFB tasks: Ball delta task = bal_de; ball theta task = bal_th; pendulum delta task = pen_de; pendulum theta task = pen_th. Horizontal bold lines indicate significant differences as revealed by the SNK post-hoc test (*p* < 0.05).

We also correlated peak frequencies during the NFB task condition with the 16 post-survey item scores. The results of these correlation analyses are presented in Figure 6. There were relatively strong positive correlations between theta peak frequency and nervousness (item 3), and negative correlations of the theta peak frequency to patience during experiment (item 6) and valued capability to influence the task through concentration and thoughts (items 13 and 15, respectively). The alpha peak frequency correlated mostly negatively, especially with general patience (item 5) and valued capability to influence the task through concentration and, to some extent, through mental calculations (items 13 and 16, respectively) during the ball task. Interestingly, during the pendulum task, this correlation was positive at fronto-central sites, when correlated with the capability to influence the task through concentration, and negative at parietal sites, when correlated with the valued capability to influence the task through relaxation. The beta peak frequency correlated strongly negatively with tiredness at the end of the experiment (item 2). In addition, there were some positive correlations between the beta peak frequency during the ball task and the valued capability to influence the task through concentration (item 13).

**Figure 6.**
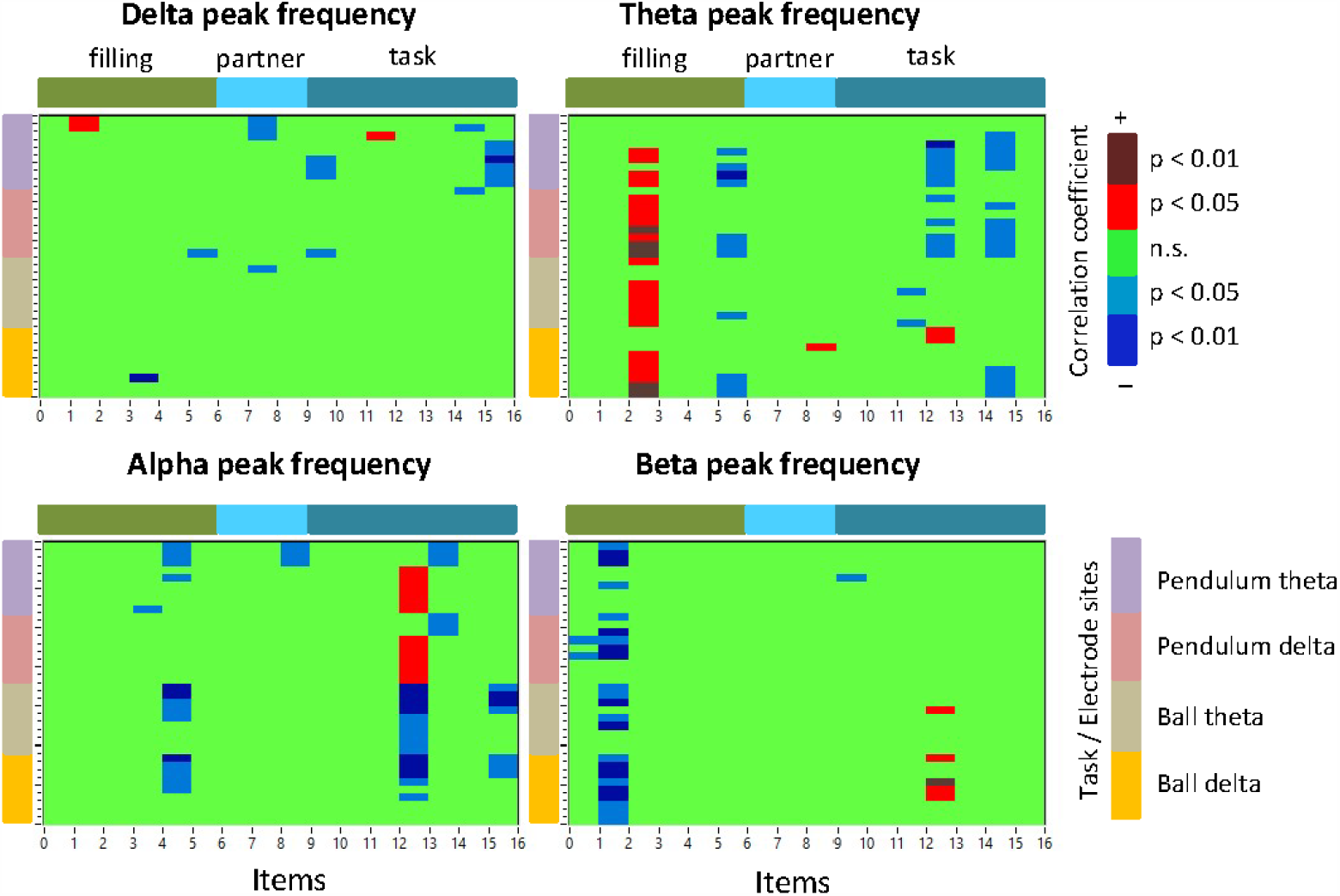
Correlations between the 16 post-survey item scores and peak frequencies in the four frequency bands. The x-axis represents the 16 post-survey item scores, which are divided into three subgroups (filing-, partner-, and task-related), as indicated by the color bars at the top of the diagrams. The y-axis represents the four tasks with nine electrode sites for each (see legend Task/Electrode sites). Electrode sites are ordered as follows: FL (frontal left), FZ (mid frontal), FR (frontal right), CL (central left), CZ (mid central), CR (central right), PL (parietal left), PZ (mid parietal), and PR (parietal right). The diagrams show only significant (negative and positive) correlation coefficients as indicated in the legend.

### Intra- and Iinter-Brain Synchrony Changes

Unlike with the power spectral indices, the *ACI* measures were determined for individual frequency bins (i.e., 2.5, 5, 10, and 20 Hz). Although the *ACI* is a symmetric measure, the topological distribution of inter-brain strengths in the two brains averaged across the nine brain sites were different. Hence, we determined the average strength for each participant and subjected them to three-way ANOVAs (*Condition* × *Antero-Posterior* × *Medio-Lateral*) similar to the analyses of spectral indices described above. The results of the within-brain coupling for the four frequency bins are presented in Table 4 and Figure 7. It can be seen that the main effect of the factor *Condition* for the intra-brain coupling strength for delta and theta frequency bins was significant in the theta pendulum task only. The intra-brain strengths in the fake condition decreased significantly compared to the rest and inverse task condition (see Fig. 7). The intra-brain alpha strengths revealed a significant main effect of *Condition* in all NFB tasks with the exception of the delta pendulum task and intra-brain beta strengths revealed a significant main effect of *Condition* in the theta ball and theta pendulum tasks. In addition, intra-brain strength in all four frequencies revealed a significant interaction *Condition* × *Antero-Posterior* among all the four NFB tasks (see Table 4 for details).

**Table 4:**
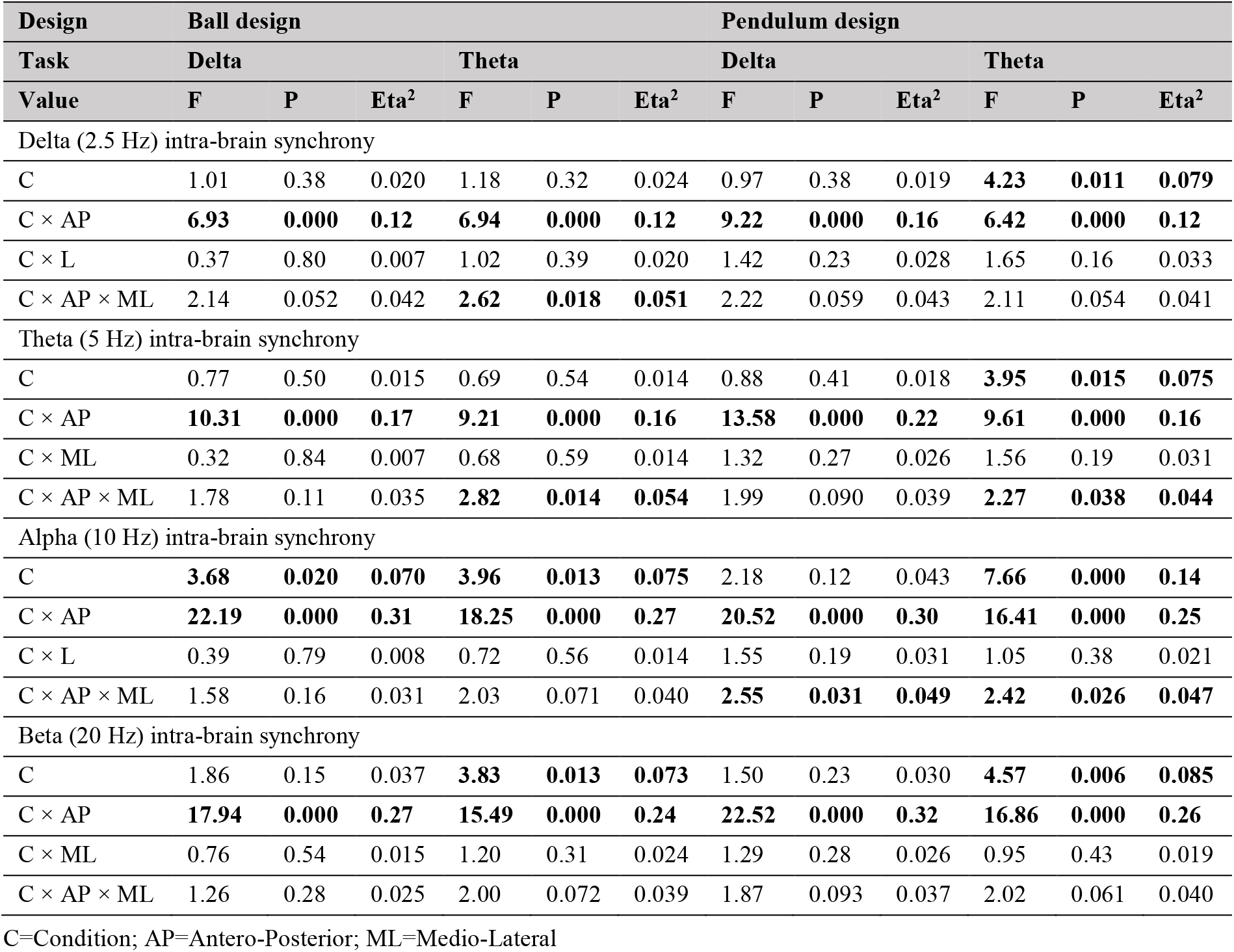
ANOVA results for the intra-brain coupling strengths across the four frequency bins (2.5, 5, 10, and 20 Hz) in the four NFB tasks

| Design | Ball design |  |  |  |  |  | Pendulum design |  |  |  |  |  |
| --- | --- | --- | --- | --- | --- | --- | --- | --- | --- | --- | --- | --- |
| Task | Delta |  |  | Theta |  |  | Delta |  |  | Theta |  |  |
| Value | F | P | Eta<sup>2</sup> | F | P | Eta<sup>2</sup> | F | P | Eta<sup>2</sup> | F | P | Eta<sup>2</sup> |
| Delta (2.5 Hz) intra-brain synchrony |  |  |  |  |  |  |  |  |  |  |  |  |
| C | 1.01 | 0.38 | 0.020 | 1.18 | 0.32 | 0.024 | 0.97 | 0.38 | 0.019 | **4.23** | **0.011** | **0.079** |
| C × AP | **6.93** | **0.000** | **0.12** | **6.94** | **0.000** | **0.12** | **9.22** | **0.000** | **0.16** | **6.42** | **0.000** | **0.12** |
| C × L | 0.37 | 0.80 | 0.007 | 1.02 | 0.39 | 0.020 | 1.42 | 0.23 | 0.028 | 1.65 | 0.16 | 0.033 |
| C × AP × ML | 2.14 | 0.052 | 0.042 | **2.62** | **0.018** | **0.051** | 2.22 | 0.059 | 0.043 | 2.11 | 0.054 | 0.041 |
| Theta (5 Hz) intra-brain synchrony |  |  |  |  |  |  |  |  |  |  |  |  |
| C | 0.77 | 0.50 | 0.015 | 0.69 | 0.54 | 0.014 | 0.88 | 0.41 | 0.018 | **3.95** | **0.015** | **0.075** |
| C × AP | **10.31** | **0.000** | **0.17** | **9.21** | **0.000** | **0.16** | **13.58** | **0.000** | **0.22** | **9.61** | **0.000** | **0.16** |
| C × ML | 0.32 | 0.84 | 0.007 | 0.68 | 0.59 | 0.014 | 1.32 | 0.27 | 0.026 | 1.56 | 0.19 | 0.031 |
| C × AP × ML | 1.78 | 0.11 | 0.035 | **2.82** | **0.014** | **0.054** | 1.99 | 0.090 | 0.039 | **2.27** | **0.038** | **0.044** |
| Alpha (10 Hz) intra-brain synchrony |  |  |  |  |  |  |  |  |  |  |  |  |
| C | **3.68** | **0.020** | **0.070** | **3.96** | **0.013** | **0.075** | 2.18 | 0.12 | 0.043 | **7.66** | **0.000** | **0.14** |
| C × AP | **22.19** | **0.000** | **0.31** | **18.25** | **0.000** | **0.27** | **20.52** | **0.000** | **0.30** | **16.41** | **0.000** | **0.25** |
| C × L | 0.39 | 0.79 | 0.008 | 0.72 | 0.56 | 0.014 | 1.55 | 0.19 | 0.031 | 1.05 | 0.38 | 0.021 |
| C × AP × ML | 1.58 | 0.16 | 0.031 | 2.03 | 0.071 | 0.040 | **2.55** | **0.031** | **0.049** | **2.42** | **0.026** | **0.047** |
| Beta (20 Hz) intra-brain synchrony |  |  |  |  |  |  |  |  |  |  |  |  |
| C | 1.86 | 0.15 | 0.037 | **3.83** | **0.013** | **0.073** | 1.50 | 0.23 | 0.030 | **4.57** | **0.006** | **0.085** |
| C × AP | **17.94** | **0.000** | **0.27** | **15.49** | **0.000** | **0.24** | **22.52** | **0.000** | **0.32** | **16.86** | **0.000** | **0.26** |
| C × ML | 0.76 | 0.54 | 0.015 | 1.20 | 0.31 | 0.024 | 1.29 | 0.28 | 0.026 | 0.95 | 0.43 | 0.019 |
| C × AP × ML | 1.26 | 0.28 | 0.025 | 2.00 | 0.072 | 0.039 | 1.87 | 0.093 | 0.037 | 2.02 | 0.061 | 0.040 |

C=Condition; AP=Antero-Posterior; ML=Medio-Lateral

**Figure 7.**
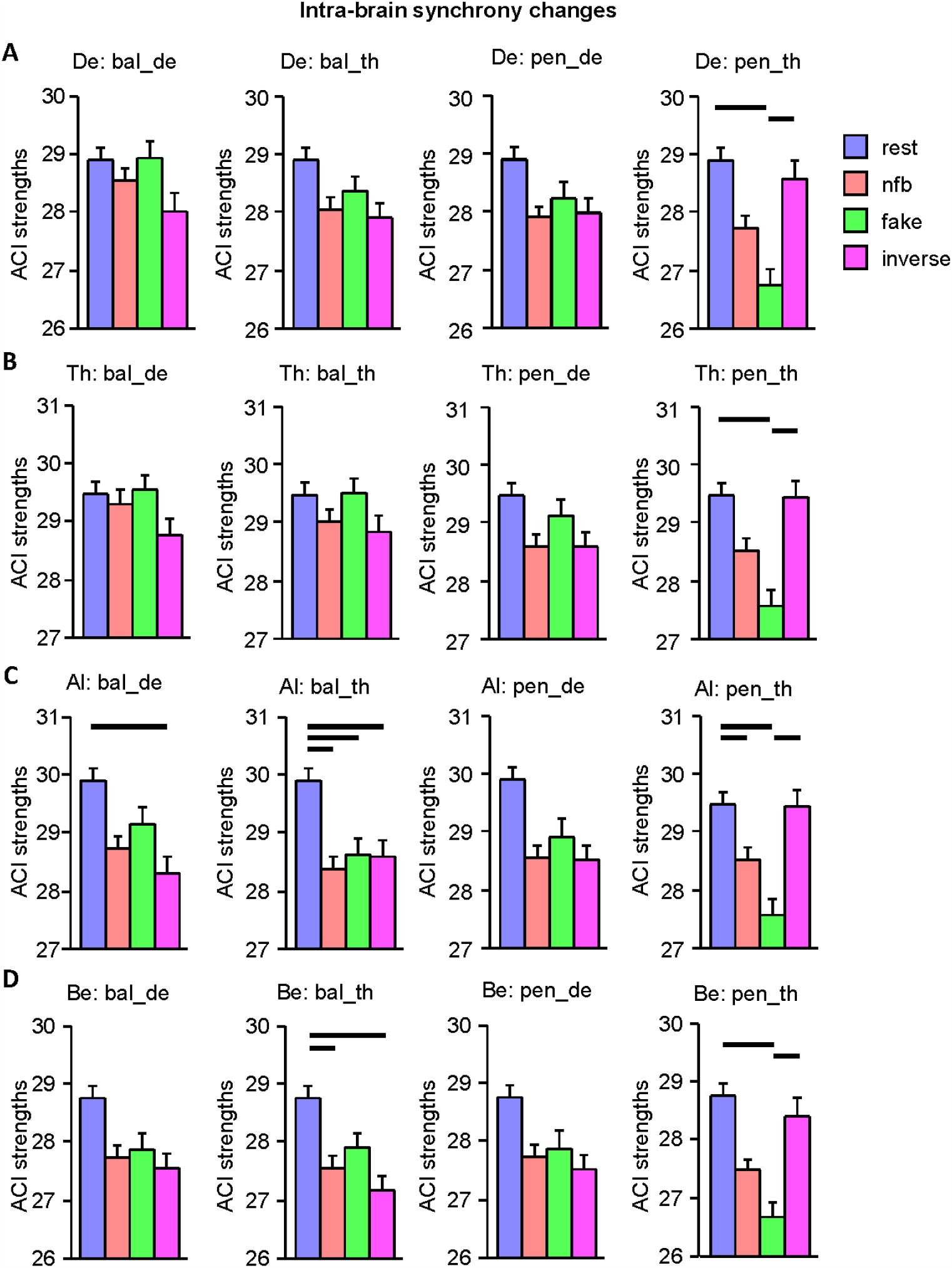
Intra-brain synchrony changes across the task conditions at the four frequency bins. (A) Delta (2.5 Hz) intra-brain synchrony across the four NFB tasks and the four task conditions. (B) Theta (5 Hz) intra-brain synchrony across the four NFB tasks and the four task conditions. (C) Alpha (10 Hz) intra-brain synchrony across the four NFB tasks and the four task conditions. (D) Beta (20 Hz) intra-brain synchrony across the four NFB tasks and the four task conditions. Bar diagrams represent means and standard errors of the *ACI* strengths across the four task conditions: resting state, normal NFB, fake, and inverse task conditions. NFB tasks: Ball delta task = bal_de; ball theta task = bal_th; pendulum delta task = pen_de; pendulum theta task = pen_th. Horizontal bold lines indicate significant differences as revealed by the SNK post-hoc test (*p* < 0.05).

As shown in Supplementary Figure S2, the intra-brain strengths in task conditions compared to the rest mainly decrease at frontal and central sites and increase at parietal sites. In the theta pendulum task, the intra-brain coupling strengths decreased in the normal and fake task conditions, but remained relatively high during the inverted task condition.

The results of the between-brain coupling for the four frequency bins are presented in Table 5 and Figure 8. Inter-brain strengths in the delta frequency (2.5 Hz) showed a significant main effect in the delta ball task and a significant interaction *Condition* × *Antero-Posterior* in the delta pendulum task. Inter-brain strengths in the three other frequencies (i.e., 5, 10, and 20 Hz) showed a significant main effect in all NFB tasks. As shown in Figure 8, the theta and beta inter-brain coupling increased significantly in the NFB task conditions compared to the rest, whereas the alpha inter-brain strengths decreased in the NFB tasks compared to the rest condition.

**Table 5:** ANOVA results for the inter-brain coupling strengths across the four frequency bins (2.5, 5, 10, and 20 Hz) in the four NFB tasks

| Design | Ball design |  |  |  |  |  | Pendulum design |  |  |  |  |  |
| --- | --- | --- | --- | --- | --- | --- | --- | --- | --- | --- | --- | --- |
| Task | Delta |  |  | Theta |  |  | Delta |  |  | Theta |  |  |
| Value | F | P | Eta<sup>2</sup> | F | P | Eta<sup>2</sup> | F | P | Eta<sup>2</sup> | F | P | Eta<sup>2</sup> |
| Delta (2.5 Hz) inter-brain synchrony |  |  |  |  |  |  |  |  |  |  |  |  |
| C | **12.43** | **0.000** | **0.20** | 2.07 | 0.11 | 0.41 | 2.76 | 0.058 | 0.053 | 1.24 | 0.29 | 0.025 |
| C × AP | 0.61 | 0.63 | 0.012 | 1.73 | 0.17 | 0.034 | **3.63** | **0.014** | **0.069** | 0.49 | 0.70 | 0.010 |
| C × ML | 0.79 | 0.52 | 0.016 | 0.85 | 0.47 | 0.017 | 0.17 | 0.93 | 0.004 | 0.92 | 0.44 | 0.018 |
| C × AP × ML | 0.93 | 0.48 | 0.019 | 0.56 | 0.79 | 0.011 | 0.66 | 0.64 | 0.013 | 0.83 | 0.54 | 0.017 |
| Theta (5 Hz) inter-brain synchrony |  |  |  |  |  |  |  |  |  |  |  |  |
| C | **10.74** | **0.000** | **0.18** | **12.85** | **0.000** | **0.21** | **4.63** | **0.006** | **0.086** | **5.10** | **0.003** | **0.094** |
| C × AP | 1.70 | 0.17 | 0.034 | 1.50 | 0.22 | 0.030 | 1.80 | 0.15 | 0.036 | 0.32 | 0.80 | 0.007 |
| C × ML | 0.62 | 0.63 | 0.012 | 1.91 | 0.13 | 0.037 | 2.08 | 0.11 | 0.041 | 0.35 | 0.82 | 0.007 |
| C × AP × ML | 1.30 | 0.26 | 0.026 | 1.64 | 0.13 | 0.032 | 1.18 | 0.32 | 0.024 | 1.72 | 0.10 | 0.034 |
| Alpha (10 Hz) inter-brain synchrony |  |  |  |  |  |  |  |  |  |  |  |  |
| C | **30.79** | **0.000** | **0.39** | **31.46** | **0.000** | **0.39** | **31.28** | **0.000** | **0.39** | **38.56** | **0.000** | **0.44** |
| C × AP | 2.21 | 0.088 | 0.043 | 1.17 | 0.32 | 0.023 | 1.62 | 0.19 | 0.032 | 1.31 | 0.28 | 0.026 |
| C × ML | 0.46 | 0.75 | 0.009 | 0.76 | 0.53 | 0.015 | 0.97 | 0.42 | 0.019 | 0.35 | 0.83 | 0.007 |
| C × AP × ML | **2.54** | **0.021** | **0.049** | 0.52 | 0.77 | 0.010 | 0.83 | 0.54 | 0.017 | 0.80 | 0.58 | 0.016 |
| Beta (20 Hz) inter-brain synchrony |  |  |  |  |  |  |  |  |  |  |  |  |
| C | **9.21** | **0.000** | **0.16** | **4.87** | **0.008** | **0.090** | **4.75** | **0.004** | **0.088** | **8.46** | **0.000** | **0.15** |
| C × AP | 0.85 | 0.48 | 0.017 | 0.72 | 0.56 | 0.014 | 0.83 | 0.48 | 0.017 | 1.52 | 0.19 | 0.030 |
| C × ML | 1.25 | 0.29 | 0.025 | 1.46 | 0.22 | 0.029 | 1.41 | 0.24 | 0.028 | 1.26 | 0.29 | 0.025 |
| C × AP × ML | 2.15 | 0.052 | 0.042 | **3.19** | **0.005** | **0.061** | 1.08 | 0.38 | 0.021 | 1.22 | 0.30 | 0.024 |

C=Condition; AP=Antero-Posterior; ML=Medio-Lateral

**Figure 8.**
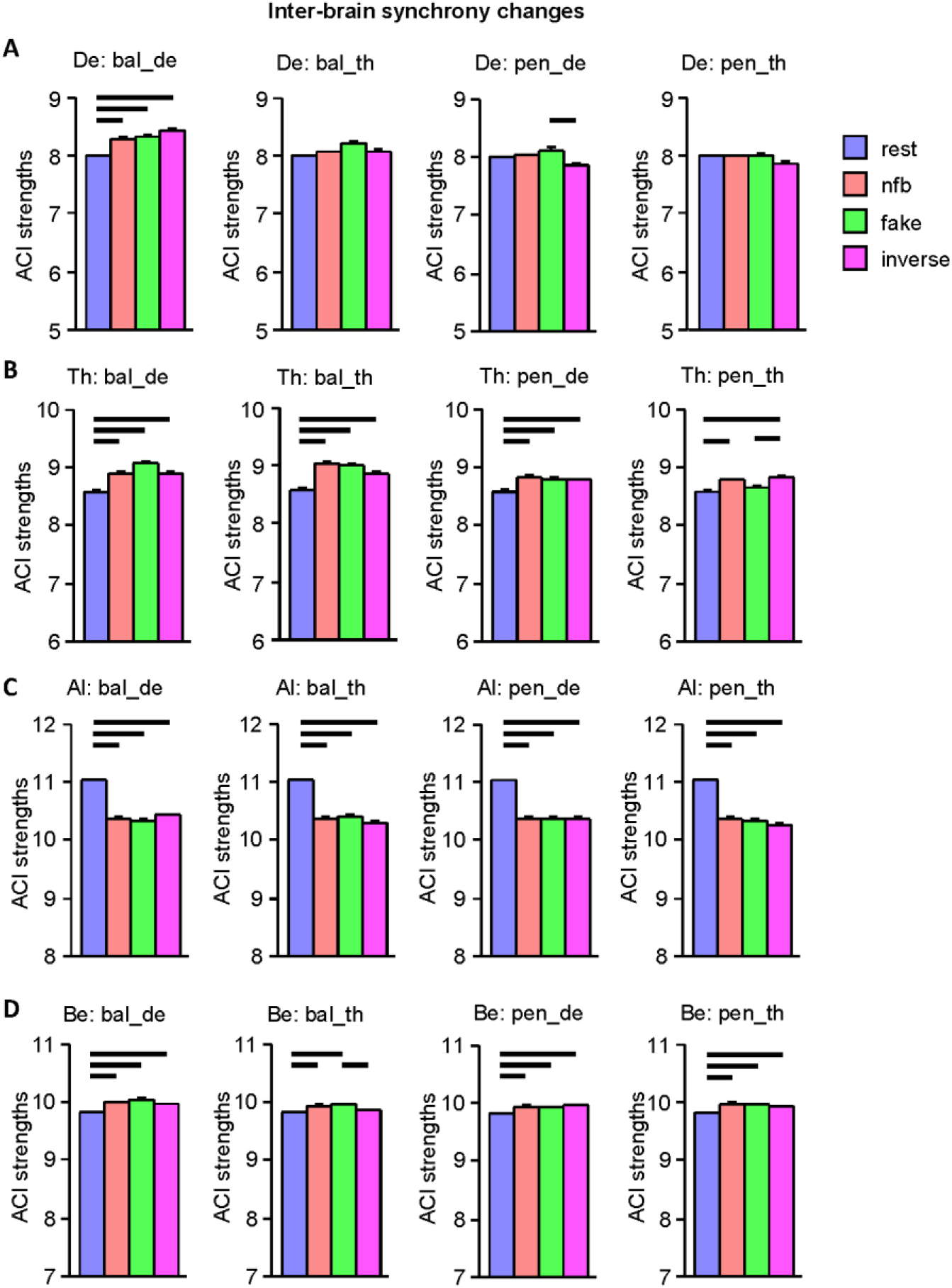
Inter-brain synchrony changes across the task conditions at the four frequency bins. (A) Delta (2.5 Hz) inter-brain synchrony across the four NFB tasks and the four task conditions. (B) Theta (5 Hz) inter-brain synchrony across the four NFB tasks and the four task conditions. (C) Alpha (10 Hz) inter-brain synchrony across the four NFB tasks and the four task conditions. (D) Beta (20 Hz) inter-brain synchrony across the four NFB tasks and the four task conditions. Bar diagrams represent means and standard errors of the *ACI* strengths across the four task conditions: resting state, normal NFB, fake, and inverse task conditions. NFB tasks: Ball delta task = bal_de; ball theta task = bal_th; pendulum delta task = pen_de; pendulum theta task = pen_th. Horizontal bold lines indicate significant differences as revealed by the SNK post-hoc test (*p* < 0.05).

Correlation analyses of intra- and inter-brain coupling strengths during the NFB task condition with the 16 post-survey item scores are presented in Figure 9. With regard to subjective feeling scores, there were negative correlations between the delta intra-brain coupling strengths and nervousness (item 3), and negative correlations between the intra-brain strengths at all frequencies and patience during the task (item 6). Beta intra-brain coupling correlated positively with the general test partner likeability (item 7), test partner sympathy (item 8), with the feeling that the synchronization did work overall well (item 10), and the feeling to have controlled the ball (and to some extent also the pendulum) during the NFB task (item 11). During the ball task, alpha intra-brain strengths also correlated positively with the feeling that the synchronization did work overall well (item 10) and, during the pendulum task, correlated negatively with the valued capability to influence the task through concentration (item 13) and through mental calculations (item 15).

**Figure 9.**
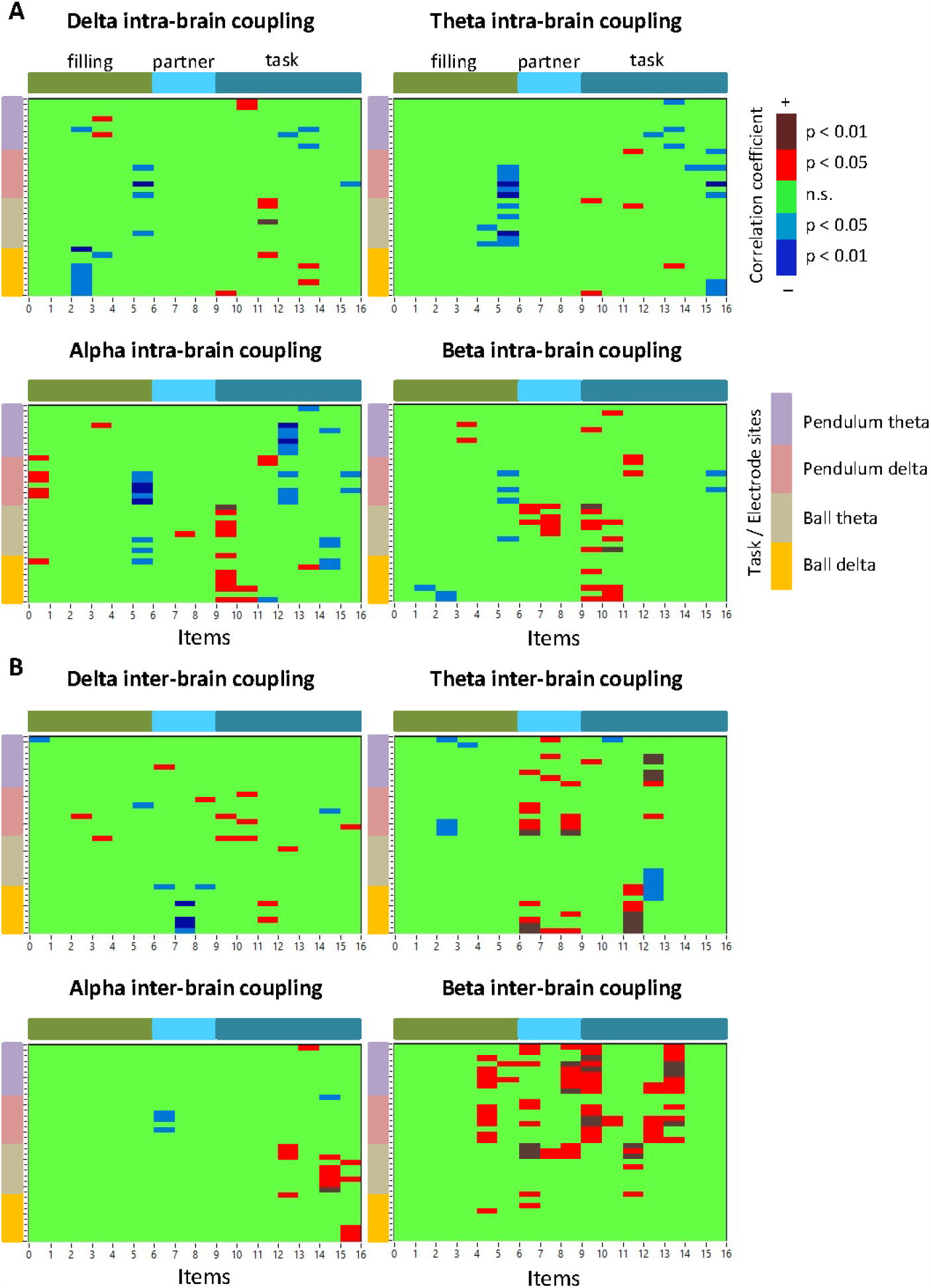
Correlations between the 16 post-survey item scores and intra- and inter-brain coupling strengths at the four frequencies. (A) Correlations between the 16 post-survey item scores and intra-brain coupling strengths. (B) Correlations between the 16 post-survey item scores and inter-brain coupling strengths. The x-axis represents the 16 post-survey item scores, which are divided in three subgroups (filing-, partner-, and task-related) as indicated by the color bars at the top of the diagrams. The y-axis represents the four tasks with nine electrode sites for each (see legend Task/Electrode sites). Electrode sites are ordered as follows: FL (frontal left), FZ (mid frontal), FR (frontal right), CL (central left), CZ (mid central), CR (central right), PL (parietal left), PZ (mid parietal), and PR (parietal right). The diagrams show only significant (negative and positive) correlation coefficients as indicated in the legend.

The inter-brain coupling strengths at theta and beta frequencies correlated significantly positively with the partner’s likability scores (items 7–9) and the feeling that the partner could control the ball better (item 12). In addition, theta inter-brain strengths correlated negatively during the ball task and positively during the pendulum task with the valued capability to influence the task through concentration (item 13), and beta inter-brain strengths correlated positively during the pendulum task with the valued capability to influence the task through concentration and relaxation (items 13 and 14, respectively). Alpha inter-brain strengths correlated significantly positively during the ball task with the valued capability to influence the task through concentration, thoughts and mental calculations (items 13, 15, and 16, respectively).

## Discussion

In the present study, we aimed to examine the neural mechanisms of interpersonal NFB. We used two experimental designs where the subjects were able to control (a) the common state without any feedback about their own contribution to this state (ball task) and (b) the common state with the feedback about their own contribution to this state (pendulum task). These states were manipulated through the enhancement of the given feedback (fake condition) and through giving an inverted feedback (inverse condition), which distracted or confused the participants. In the ball task, the inverse feedback led to the appearance of mostly overlapped balls falsely indicating a good performance whereas, in the pendulum task with inverse feedback, the pendulums were swinging most the time in antiphase, falsely indicating a bad performance. Thus, whereas, in the pendulum task, the participants were really frustrated by the negative feedback, in the ball task, the participants could not understand why they were so good at times, and were, thus, frustrated or disoriented by the overwhelming positive feedback.

EEG analyses revealed strong differences between the resting state and the NFB task conditions, but the differences between the NFB task conditions themselves were rather moderate or mostly absent. PSD analyses showed a strong decrease of PSD in all frequency bands in the NFB task conditions compared to the rest condition. Regarding the NFB task conditions themselves, there was only a significant decrease of beta PSD in the inverse task condition compared to the normal NFB condition. This indicates that the NFB tasks led to the common desynchronization of all EEG rhythms compared to the rest. Such a decrease in spectral power (at least in the frequency ranges of 8–20 Hz and 35–45 Hz) was also observed in the study on collective NFB mentioned above (Kovacevic et al., 2015). Also, Ros et al. (2013) reported an amplitude or spectral power reduction in the theta, alpha, and beta bands during the NFB task. As suggested by the authors and other investigators (Fan et al., 2007; Friese et al., 2016), such broadband spectral power attenuation or desynchronization could be a sign of alerting and selective attention.

Interestingly, PSD at all frequencies showed significant positive correlations with tiredness after the experiment and nervousness (except delta PSD), which were especially strong at fronto-central sites. Note that these brain sites were used for the feedback calculation and that the tiredness before the experiment did not show any significant correlations. Moreover, the PSD (also in all frequency bands) correlated significantly negatively with the sympathy for the test partner and also with the valued capability to influence the task. Interestingly, the latter were also found at the fronto-central sites. This indicates that a too higher PSD led to tiredness and nervousness and, at the same time, lower PSD seems to positively influence the feeling of good performance or task controllability and the test partner’s sympathy. There is evidence that tiredness is especially associated with low-frequency oscillations (Wyczesany et al., 2008), and it has been suggested that alpha and beta power are related to cognitive and emotional control (Ray and Cole, 1985). Also, in the study on collective NFB (Kovacevic et al., 2015), the decrease in broadband spectral power was associated with the ability of the participants to maintain the desired state.

Furthermore, we found that the spectral peak frequency decreases in the delta frequency range (only in the pendulum task) and especially in the theta frequency range. Importantly, the theta peak frequency converges to the task frequency of 5 Hz. Alpha and beta peak frequencies showed only significant interactions of the factor Condition with electrode sites, indicating different changes with regard to the different brain regions, whereby this was the lateral axis in the alpha frequency band and the antero-posterior axis in the beta frequency band. It should be noted that the alpha and the beta peak frequencies rather increased compared to the rest. Most importantly, these frequencies (i.e., alpha and beta) also showed changes within the test pairs, namely, the differences in peak frequencies within the pairs increase compared to the rest whereas, in the NFB-task frequencies (i.e., delta and theta), there were no differences within the pairs. It seems that the adjustment of the task rhythms (i.e., delta and theta) happens due to, or is accompanied by, the divergence of the fast rhythms (alpha and beta) within the test pairs. Interestingly, the theta peak frequency correlated significantly positively with nervousness, especially at fronto-central sites, and significantly negatively with patience during the experiment and also with the valued capability to influence the task through concentration and thoughts. The beta peak frequency correlated significantly negatively with tiredness after the experiment. This only confirms the suggestion that the decrease in theta peak frequency (and partly also in the delta peak frequency) and increase in the beta peak frequency (at least at the fronto-central brain regions) could be beneficial for the task and associated with less nervousness.

Intra-brain synchrony either shows no changes during the tasks or rather decreases compared to the resting state. Inter-brain synchrony during the tasks increases compared to rest in the theta and beta frequency ranges and partly also in the delta frequency (e.g., during the delta ball task). Inter-brain synchrony at the alpha frequency decreases significantly during the task conditions compared to rest. The alpha and beta frequencies were not manipulated during the task, but a significant decrease of inter-brain synchrony in the former case and an increase in the latter indicate that these frequencies are also drawn into passion. It simply shows that if the participants concentrate on a certain rhythm and change it accordingly, the whole frequency spectrum changes. It is interesting to note that some accompanying frequencies desynchronize in relation to the partner (e.g., alpha frequency) and other frequencies synchronize (e.g., beta frequency). In our opinion, this is an interesting phenomenon that needs further investigation to be better understood. Especially interesting is the strongly significant decrease in alpha inter-brain synchronization in the NFB task conditions compared to the rest condition. In principle, inter-brain synchronization during the rest could be regarded as a spurious synchronization, but the fact that during the NFB task (regardless of the task design or task frequency), it dropped drastically indicates that it is then more spurious than in rest. In our opinion, alpha oscillations during rest are relatively stable in both participants, whereas during the task they move apart or desynchronize in the test pair. This view is in line with the significant increase of peak frequency differences within the test pairs during the task (see Figure 5). Further, it should be noted that theta and beta inter-brain coupling strengths correlated mostly significantly positively with partner- and task-oriented post-survey scores. This indicates that inter-brain synchrony in these frequency bands (i.e., theta and beta), but also partly in the delta and alpha frequency bands, are relatively strongly related to the test partner’s likability and valued capability to influence the task or task frequency.

There is neurophysiological evidence that at least two large-scale neural networks that represent the self and others play a crucial role in self-related processing and social cognition (Uddin et al., 2007). These are frontoparietal mirror-neuron areas connecting the physical self and others through motor-simulation mechanisms and cortical midline structures engaging in processing information about the self and others in more abstract, evaluative terms (Uddin et al., 2007), called also a midline self-representation core (Ioannides, 2018). Interestingly, these networks are also activated for NFB interventions that are highly personalized and that attempt to alter the neural self-representation (Ioannides, 2018). The mirror neuron system, which shows similar activation during perception and execution of action (Schmidt et al., 2011) – a process termed common coding of perception and action (Novembre et al., 2014; Sebanz and Knoblich, 2009) – implies that the processing of other people’s actions relies on the simulation of those actions in one’s own motor system (Wolpert et al., 2003; Wolpert and Ghahramani, 2000). In our NFB tasks, the participants have to mirror or predict the internal state of others and simulate their activity in the respective partner-oriented way. It has been suggested that, during an interpersonal interaction, each subject has their own forward model and corresponding neural representation of this, whereby these representations are adjusted in time (Sänger et al., 2011). Moreover, they set boundary conditions for the agents to behave as a superordinate system or as a whole (cf. (Müller et al., 2018a)). We have shown in this study that, during the interpersonal NFB task, there are specific brain dynamics both within and between the brains, which is also related to behavioral post-survey item scores. Nevertheless, further sophisticated research is needed to deepen and extend our understanding of these highly interesting and complex phenomena.

## Limitations

The present experiment has limitations and leaves room for questions to be addressed in future research. First, we found strong differences in neural activity and synchrony during the task compared to the resting state. Using more fine-grained NFB task manipulations would lead to a more differentiated representation of NFB activity and reinforce our understanding of both NFB mechanisms and inter-brain or hyper-brain interaction. Second, our analyses with regard to intra- and inter-brain coupling were limited to the *ACI* measure reflecting the in-phase synchrony, which was important with regard to the experimental designs. Nevertheless, other types of coupling (e.g., directional or nonlinear coupling as well as multivariate coupling) are likely to provide further information about the synchronized states during interpersonal NFB.

## Conclusion

Our results show that the participants were able to increase inter-brain synchrony by using NFB information, especially when the inter-brain coupling was fed back at the theta frequency. Apart from the inter-brain coupling other oscillatory activities (e.g., PSD, peak amplitude and peak frequency, intra-brain coupling) changed during the task compared to the rest. Moreover, all the measures showed specific correlations to the subjective post-survey item scores indicating specific relations of neural oscillatory activity to the subjective feeling and appraisal. Finally, this study shows that hyperscanning with IBS feedback seems to be an important tool to examine inter-brain oscillatory coupling and performance that provides important information about neural mechanisms of social interaction and collective behavior.

## Acknowledgments

This research was supported by the Max Planck Society. The authors are grateful to Oksana Berhe, Vivien Choupurian, Shiva Motlagh, Katrin Müller, and Alireza Tarikhi for technical assistance and for carrying out the experimental part of the study. The authors thank Christel Fraser for language assistance, and Berndt Wischnewski and Ioanna Michopoulou for software assistance.

## Competing interests

The authors declare no competing interests.

## Notes

### Competing Interest Statement

The authors have declared no competing interest.

