## Supplementary material for "Interacting brains coming in sync through their minds: An inter-brain neurofeedback study": Mueller etal_supp_BioRxiv.pdf

Supplementary Table S1: Post-survey items

| Item category /number | Item | Answer |
|---|---|---|
| Filling | | |
| 1 | How tired were you at the beginning of the experiment? | 1=not tired, 5=very tired |
| 2 | How tired were you at the end of the experiment? | 1=not tired, 5=very tired |
| 3 | How nervous do you feel today? | 1=not nervous, 5=very nervous |
| 4 | Do you have an important appointment today? | 1=no important, 5=very important |
| 5 | How patient do you consider yourself to be in general? | 1=not patient, 5=very patient |
| 6 | How patient did you think you were today during the neurofeedback task? | 1=not patient, 5=very patient |
| Partner-oriented | | |
| 7 | Reysen Likeability Scale (averaged across 11 items) | 1=disagree very strongly, 7=agree very strongly |
| 8 | How sympathetic was your test partner to you? | 1=not at all, 5=very sympathetic |
| 9 | How similar do you rate the personality of your test partner compared to your own personality? | 1=completely different, 5=identical |
| Valued capability to influence the task | | |
| 10 | How well did the synchronization work overall? | 1=bad, 5=perfect |
| 11 | How much did you overall have the feeling of being able to control the pendulum/the ball? | 1=bad, 5=perfect |
| 12 | Who controlled the pendulum/the ball more - you or your partner? | 1=mostly me, 5=mostly my partner |
| 13 | How much did you control the pendulum/the ball using the following strategies: *concentration*? | 1=not at all, 5=very strongly |
| 14 | How much did you control the pendulum/the ball using the following strategies: *relaxation*? | 1=not at all, 5=very strongly |
| 15 | How much did you control the pendulum/the ball using the following strategies: *generating thoughts*? | 1=not at all, 5=very strongly |
| 16 | How much did you control the pendulum/the ball using the following strategies: *mental calculation*? | 1=not at all, 5=very strongly |

Reysen Likeability Scale (Reysen, 2005) includes 11 items with 7 answer possibilities: 1) disagree very strongly, 2) disagree strongly, 3) disagree, 4) neutral, 5) agree, 6) agree strongly, and 7) agree very strongly. The scores were averaged across these items. The items reflecting the valued capability to influence the task (10-16) were always asked separately for ball and pendulum tasks, respectively.

Supplementary Table S2: ANOVA results for the peak amplitude across the four frequency bands in the four NFB tasks

| **Design** | **Ball design** | | | | | | **Pendulum design** | | | | | |
|---|---|---|---|---|---|---|---|---|---|---|---|---|
| **Task** | **Delta** | | | **Theta** | | | **Delta** | | | **Theta** | | |
| **Value** | **F** | **P** | **Eta$^2$** | **F** | **P** | **Eta$^2$** | **F** | **P** | **Eta$^2$** | **F** | **P** | **Eta$^2$** |
| Delta peak amplitude | | | | | | | | | | | | |
| C | **5.79** | **0.005** | **0.11** | **5.94** | **0.005** | **0.11** | 2.77 | 0.072 | 0.053 | 3.05 | 0.054 | 0.059 |
| C × AP | **2.84** | **0.044** | **0.055** | 2.06 | 0.10 | 0.040 | 1.97 | 0.12 | 0.039 | 2.03 | 0.11 | 0.040 |
| C × L | 1.44 | 0.22 | 0.029 | 1.14 | 0.34 | 0.023 | 0.45 | 0.81 | 0.009 | 0.73 | 0.58 | 0.015 |
| C × AP × ML | 0.76 | 0.63 | 0.015 | 1.01 | 0.43 | 0.020 | 1.47 | 0.18 | 0.029 | **2.57** | **0.011** | **0.050** |
| Theta peak amplitude | | | | | | | | | | | | |
| C | **32.51** | **0.000** | **0.40** | **32.76** | **0.000** | **0.40** | **22.35** | **0.000** | **0.31** | **29.08** | **0.000** | **0.37** |
| C × AP | **5.38** | **0.001** | **0.099** | **5.75** | **0.001** | **0.11** | **3.74** | **0.011** | **0.071** | **5.43** | **0.002** | **0.10** |
| C × ML | **3.46** | **0.010** | **0.066** | **2.66** | **0.033** | **0.051** | 0.74 | 0.58 | 0.015 | 1.53 | 0.20 | 0.030 |
| C × AP × ML | 0.44 | 0.87 | 0.009 | 0.52 | 0.79 | 0.010 | 1.00 | 0.42 | 0.020 | **3.04** | **0.004** | **0.058** |
| Alpha peak amplitude | | | | | | | | | | | | |
| C | **115.55** | **0.000** | **0.70** | **101.87** | **0.000** | **0.68** | **78.50** | **0.000** | **0.62** | **80.40** | **0.000** | **0.62** |
| C × AP | **17.46** | **0.000** | **0.26** | **19.36** | **0.000** | **0.28** | **17.03** | **0.000** | **0.26** | **17.19** | **0.000** | **0.26** |
| C × ML | 1.02 | 0.40 | 0.020 | 0.52 | 0.71 | 0.010 | 1.35 | 0.25 | 0.027 | 0.34 | 0.84 | 0.007 |
| C × AP × ML | 1.56 | 0.16 | 0.031 | 1.69 | 0.13 | 0.033 | 0.76 | 0.58 | 0.015 | 1.88 | 0.078 | 0.037 |
| Beta peak amplitude | | | | | | | | | | | | |
| C | **27.59** | **0.000** | **0.36** | **33.61** | **0.000** | **0.41** | **20.36** | **0.000** | **0.29** | **19.48** | **0.000** | **0.28** |
| C × AP | **6.45** | **0.000** | **0.12** | **7.85** | **0.000** | **0.14** | **9.72** | **0.000** | **0.17** | **8.18** | **0.001** | **0.14** |
| C × ML | 1.18 | 0.32 | 0.023 | 0.41 | 0.82 | 0.008 | 1.30 | 0.27 | 0.026 | 0.97 | 0.43 | 0.019 |
| C × AP × ML | 0.43 | 0.90 | 0.009 | 1.30 | 0.26 | 0.026 | 1.29 | 0.26 | 0.026 | **2.50** | **0.013** | **0.049** |

C=Condition; AP=Antero-Posterior; ML=Medio-Lateral

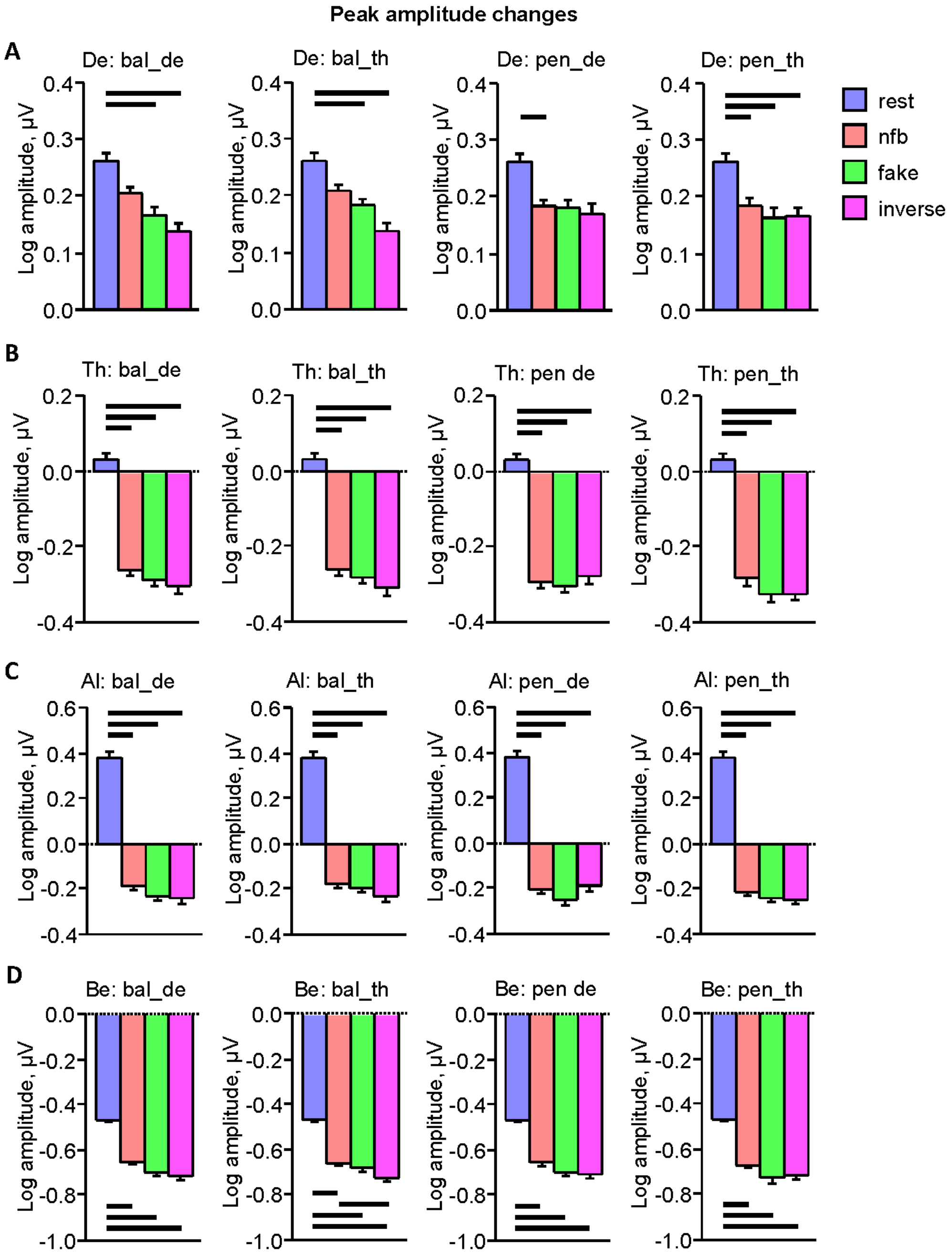


**Supplementary Figure S1.** Peak amplitude changes across the task conditions in the four frequency bands. (A) Delta peak amplitude across the four NFB tasks and the four task conditions. (B) Theta peak amplitude across the four NFB tasks and the four task conditions. (C) Alpha peak amplitude across the four NFB tasks and the four task conditions. (D) Beta peak amplitude across the four NFB tasks and the four task conditions. Bar diagrams represent peak amplitude means and standard errors across the four task conditions: resting state, normal NFB, fake, and inverse task conditions. NFB tasks: Ball delta task = bal_de; ball theta task = bal_th; pendulum delta task = pen_de; pendulum theta task = pen_th. Horizontal bold lines indicate significant differences as revealed by the SNK post-hoc test ($p < 0.05$).

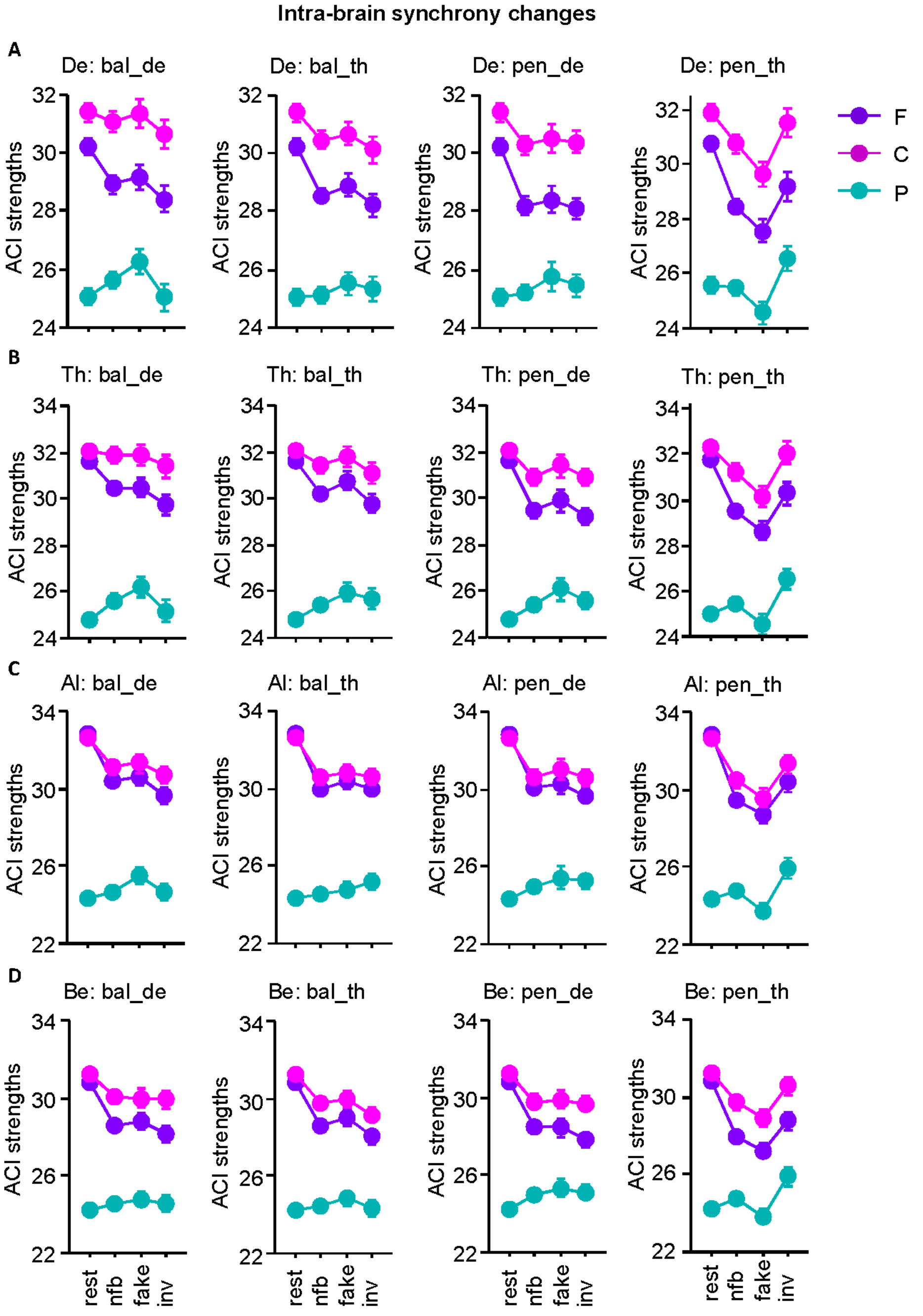


**Supplementary Figure S2.** Intra-brain synchrony changes along the antero-posterior axis across the task conditions at the four frequency bins. (A) Delta (2.5 Hz) intra-brain synchrony at fronto-parietal sites across the four NFB tasks and the four task conditions. (B) Theta (5 Hz) intra-brain synchrony at fronto-parietal sites across the four NFB tasks and the four task conditions. (C) Alpha (10 Hz) intra-brain synchrony at fronto-parietal sites across the four NFB tasks and the four task conditions. (D) Beta (20 Hz) intra-brain synchrony at fronto-parietal sites across the four NFB tasks and the four task conditions. Line diagrams with means and standard errors of the *ACI* strengths represent the *Condition* by *Antero-Posterior* interactions. Task conditions: resting state, normal NFB, fake, and inverse task conditions. NFB tasks: Ball delta task = bal_de; ball theta task = bal_th; pendulum delta task = pen_de; pendulum theta task = pen_th. Antero-posterior sites: F = frontal; C = central; P = parietal.

### Supplementary movies

File names:
NF_ball_nfb_delta.mp4
NF_ball_nfb_theta.mp4
NF_ball_fake_delta.mp4
NF_ball_fake_theta.mp4
NF_ball_inverse_delta.mp4
NF_ball_inverse_theta.mp4
NF_pendulum_nfb_delta.mp4
NF_pendulum_nfb_theta.mp4
NF_pendulum_fake_delta.mp4
NF_pendulum_fake_theta.mp4
NF_pendulum_inverse_delta.mp4
NF_pendulum_inverse_theta.mp4

File format: mp4

Description of movies: These movies show the simulation data of the different NFB tasks with two balls or pendula as they were seen on the screen during the experiment. Please note that the diagrams on the bottom, indicating changes in *ACI* (Absolute Coupling Index) in the ball task or phases in the pendulum task, were not seen during the experiment and were added for simulation and visualization purposes.
